# Coordinated yet asymmetric striatal neuromodulatory dynamics encode associative learning

**DOI:** 10.64898/2026.03.17.712528

**Authors:** Min Jung Kim, Yang Yang, Pemantha Lakraj Gamage, Tabitha Haun, Yize Wu, Dayelin Navarro, Nan Li

**Affiliations:** Advanced Imaging Research Center, University of Texas, Southwestern Medical Center, 5323 Harry Hines Blvd., Dallas, TX, 75390; Department of Neuroscience, University of Texas, Southwestern Medical Center, 5323 Harry Hines Blvd., Dallas, TX, 75390; Peter O’Donnell Jr. Brain Institute, University of Texas, Southwestern Medical Center, 5323 Harry Hines Blvd., Dallas, TX, 75390; Department of Mathematics and Statistics, University of North Carolina at Greensboro, 1400 Spring Garden Street, Greensboro, NC 27412

**Author notes:** Correspondence and request for materials should be addressed to NL.

## Abstract

The striatum integrates dopamine and acetylcholine to support learning and behavioral flexibility, yet how these neuromodulators coordinate during associative learning remains unclear. Using longitudinal dual-color fiber photometry in mice, we simultaneously tracked both signals in the anterior dorsolateral striatum throughout Pavlovian conditioning. We find that learning recruits distinct forms of neuromodulatory plasticity: dopamine maintains rapid, contingency-dependent cue responses, whereas acetylcholine undergoes broader reorganization across training. Despite these differences, trial-by-trial covariance between the two signals converges onto a compact low-dimensional structure that reliably captures learned behavioral state and distinguishes associative learning from sensory exposure alone. Granger causality analyses further reveal a directional asymmetry, present in both paired and unpaired mice, whereby dopamine reliably predicts subsequent acetylcholine fluctuations, with only weak reciprocal prediction. Together, our results identify a low-dimensional organizational principle for dopamine–acetylcholine interactions, and suggest that dopamine acts as a leading temporal signal that shapes cholinergic dynamics during striatal learning.

## Introduction

The striatum is a key nexus for transforming cortical inputs into flexible motor actions, a process critically dependent on the plasticity of spiny projection neurons (SPNs)^1-4^. This plasticity is governed by a powerful yet incompletely understood synergy between dopamine (DA) and acetylcholine (ACh)^5-7^. DA provides a canonical reward prediction error (RPE) readout^8-11^, yet context-specific deviations and circuit heterogeneity persist^12-16^; it also drives corticostriatal potentiation through D1/D2 receptor signaling^1,2^, whereas ACh acts as a rapid gain controller, modulating SPN excitability and synaptic input selection via muscarinic and nicotinic receptors^17,18^. At the local circuit level, DA terminals can suppress cholinergic interneurons (CINs) activity via D2 receptors ^19^, while synchronized CINs can conversely drive DA release through nicotinic receptors ^20^. Despite the well-established importance of this dual neuromodulatory system^21^, how DA and ACh coordinate their activity to jointly organize striatal computation during adaptive learning remains unsolved ^22^.

Earlier studies have provided substantial insight into DA-ACh interactions through a variety of experimental strategies. Loss-of-function experiments, in which one input was ablated while monitoring the other, suggested a marked degree of autonomy between the two systems^23-25^. Transient coactivation analyses, such as phasic DA bursts coinciding with ACh pauses, further identified short-timescale correspondences between the two signals, but offered limited understanding of their behavioral relevance or the directional temporal dependencies that underlie learning^17,25,26^. Even studies combining electrophysiological recordings of CINs with dopaminergic measurements (e.g., fast-scan cyclic voltammetry, single-color photometry) have typically operated across separate experimental contexts or brief recording windows^14,26^, leaving behaviorally embedded DA-ACh interactions difficult to resolve directly.

A further limitation of prior studies is the widespread reliance on trial-averaged responses. Although averaging enhances event-locked signals, it obscures temporal heterogeneity, cross-channel covariance, and dynamic features critical for encoding complex learning state^27-29^. Because neuromodulator release in the striatum is inherently variable and ensemble-based^18,24^, resolving their joint function requires analyses that capture trial-by-trial interactions rather than reducing each neurotransmitter to an independent scalar measure ^28-30^. Consequently, it remains unclear how DA and ACh evolve together across learning trials, whether they encode behavioral states jointly or independently, and whether their interaction has any directional structure.

To address these challenges, we performed longitudinal dual-color fiber-photometry to simultaneously track DA and ACh activity in the anterior dorsolateral striatum (aDLS) across the full trajectory of cue–reward associative learning. This approach provided real-time, trial-resolved, behaviorally aligned measurements of both neuromodulators throughout acquisition, together with validation of signal integrity and channel independence ^14,31,32^. We combined functional regression, dimensionality reduction^27,28^, and Granger causality analyses to determine how the interactions between DA and ACh evolve across task epochs and spontaneous behavior.

We found that associative learning stabilized fast cue-evoked DA transients, while broadly reorganizing ACh dynamics, and restructured spontaneous licking into more stereotyped bouts with selectively enhanced cholinergic encoding of lick-burst vigor. Across trials, the dominant patterns of joint DA–ACh covariance formed a low-dimensional structure that robustly encoded trial-wise learned state. While ACh motifs captured broader population-level state transitions, DA motifs contributed more selectively to learned-state discrimination. Furthermore, Granger causality revealed a sustained directional asymmetry with DA reliably predicting ACh fluctuations far more strongly than the reverse. Together, these results identify a hierarchical neuromodulatory organization in which coordinated low-dimensional DA–ACh dynamics encode learned behavioral state, with DA acting as a temporally preceding signal that scaffolds ACh activity, thereby reframing these neuromodulators not as independent channels but as a coupled neuromodulatory system for striatal computation that supports adaptive learning^1,3,24^.

## Result

### Lick-pattern clustering tracks acquisition of cue responding

To provide a context for studying neuromodulatory signaling dynamics during learning, mice were trained on a Pavlovian cue-reward association task. G-protein-coupled receptor activation-based (GRAB) sensors were used for simultaneously tracking striatal dopamine (DA) and acetylcholine (ACh) release with precise behavioral alignment.

To establish baseline responses, mice first completed identical cue-only and random-reward sessions before being assigned to training groups. They then underwent daily Pavlovian cue-reward conditioning, receiving either paired (cue followed by contingent reward) or unpaired (no contingency) training across 15 sessions (Fig. 1a). In the paired group, lick responses to the cue became progressively stereotyped over sessions (Fig. 1b).

**Figure 1.**
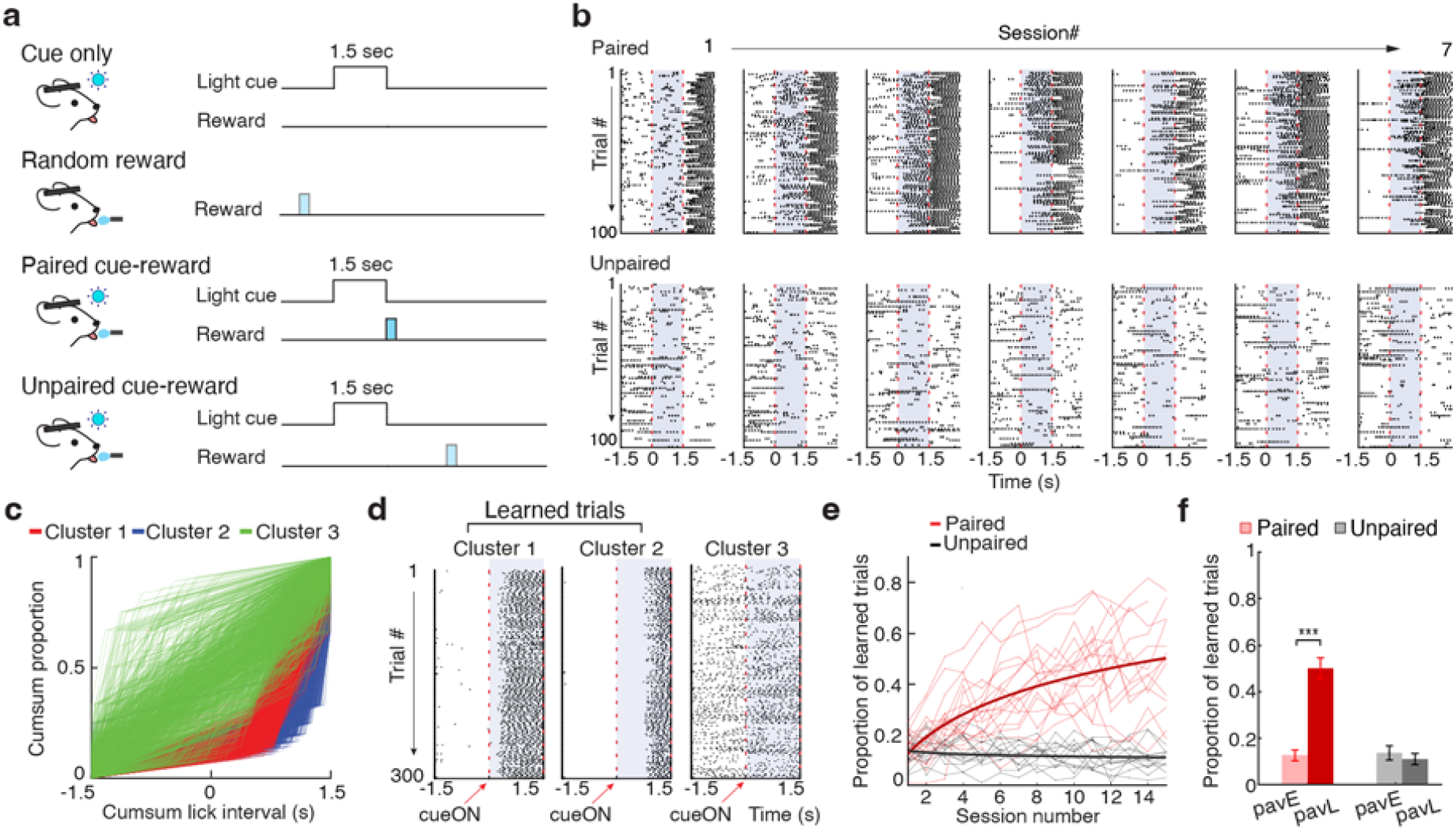
Paired Pavlovian conditioning drives the progressive emergence of cue-locked anticipatory licking. a. Schematic of behavioral paradigms including visual representations of cue-only, random reward, and paired versus unpaired cue-reward contingencies. **b**. Evolution of licking behavior across sessions is shown through representative lick rasters from a single mouse over seven training sessions with 100 trials per session. The paired mouse (top) shows the progressive emergence of anticipatory licking, whereas the unpaired mouse (bottom) shows diffuse and non-cue-locked licking. Shaded blue areas indicate the 1.5-second cue period. **c**. Quantitative trial classification based on the distribution of three distinct lick clusters identified across the population. Classification was determined by the cumulative proportion of inter-lick intervals during the pre-cue and cue periods. **d**. Cluster-specific lick rasters representing licking patterns with 300 trials per cluster aligned to cue onset marked by the red dashed line. Trials assigned to Clusters 1 and 2 are categorized as learned trials, while Cluster 3 and trials with no licking are categorized as unlearned trials. **e**. Learning trajectories across cohorts demonstrating the acquisition of the learned licking response modeled using a binomial generalized linear mixed model (GLMM). Paired mice (red, n=20) show a significant log-linear increase in learned trial proportion compared to the flat trajectory observed in unpaired mice (black, n=13). **f**. Summary of behavioral acquisition showing the marginal mean proportion of learned trials in early (pavE) versus late (pavL) sessions. Error bars indicate 95% confidence intervals. Significance markers denote Wilcoxon signed-rank tests where *** *p* < 0.001.

To quantify trial-level variation in learned responses, we clustered licking behavior within a 1.5 s window around cue onset based on the cumulative distribution of inter-lick intervals (Fig. 1c). This analysis identified three distinct clusters of trials: two characterized by minimal pre-cue licking followed by tightly time-locked licking after cue onset, classified as learned trials (tcL), and a third cluster showing disorganized or premature licking, which was grouped with no-lick trials and classified as not learned (tcNL) (Fig. 1d).

Across training, the proportion of tcL trials in each session increased robustly in paired mice (n = 20) but remained low in unpaired controls (n = 13; Fig. 1e). Importantly, this increase reflected a probabilistic shift in trial composition rather than a uniform transition of each animal to exclusively learned responding. The within-session mixture of tcL and tcNL trials, persisting across stages of training and varying in composition across individual animals, enabled trial-resolved behavioral labeling necessary for subsequent trial-by-trial neuromodulatory decoding analyses.

To characterize learning trajectories with higher temporal resolution, we modeled the probability of a learned trial across all 15 sessions using a binomial generalized linear mixed-effects model (GLMM). Model comparison strongly favored a log-linear acquisition function over a linear alternative (ΔAkaike information criterion, ΔAIC = 156), consistent with rapid early learning followed by a gradual asymptote. The selected model revealed a significant effect of session (logSess, β = 0.696, *p* = 5.41 × 10^-144^), reflecting a strong overall increase in conditioned responding. While the main effect of group was non-significant (β = 0.104, *p* = 0.535), indicating equivalent behavioral baselines, the Group × Session interaction was highly significant (β = −0.786, *p* = 2.29 × 10^-81^). To evaluate within-group behavioral changes, we compared the proportion of learned trials in early (Session 1, PavE) vs. late (Session 15, PavL) training phases (Fig. 1f). For the paired group, a Wilcoxon signed-rank test revealed a significant increase in conditioned responding (*p* < 0.001), whereas no significant change was observed for the unpaired group (*p* = 0.412).

This demonstrates that the rate and shape of acquisition diverged sharply between cohorts: paired mice exhibited a steep, logarithmic increase in learned trials, whereas unpaired mice failed to show systematic improvement. These results establish that conditioned licking emerges progressively and is contingent on the cue–reward association.

### Session-averaged neuromodulator responses diverge by learning stimuli

To monitor neuromodulatory dynamics during learning, we expressed green fluorescent GRAB-ACh indicator gACh4h^33^ and red fluorescent GRAB-DA indicator rDA2m^34^ in the anterior dorsolateral striatum (aDLS; Fig. 2a) and performed longitudinal dual-color fiber photometry across all training sessions. Signal integrity was validated by examining the relationship between the active gACh4h 470 nm channel and its 405 nm isosbestic reference, which exhibited reciprocal dynamics across recordings with the reference channel fluctuating at much smaller amplitude (Fig. S1), indicating that the active-channel signal predominantly reflects sensor-derived ACh activity rather than nonspecific artifact. In addition, cross-channel independence and neurochemical specificity were confirmed using a non-binding mutant dopamine sensor^35^ (Fig. S2). Representative trial-wise and session-averaged traces of ACh and DA signals aligned to cue and reward events showed the emergence of structured, event-specific dynamics from early (PavE) to late (PavL) Pavlovian conditioning (Fig. 2b).

**Figure 2.**
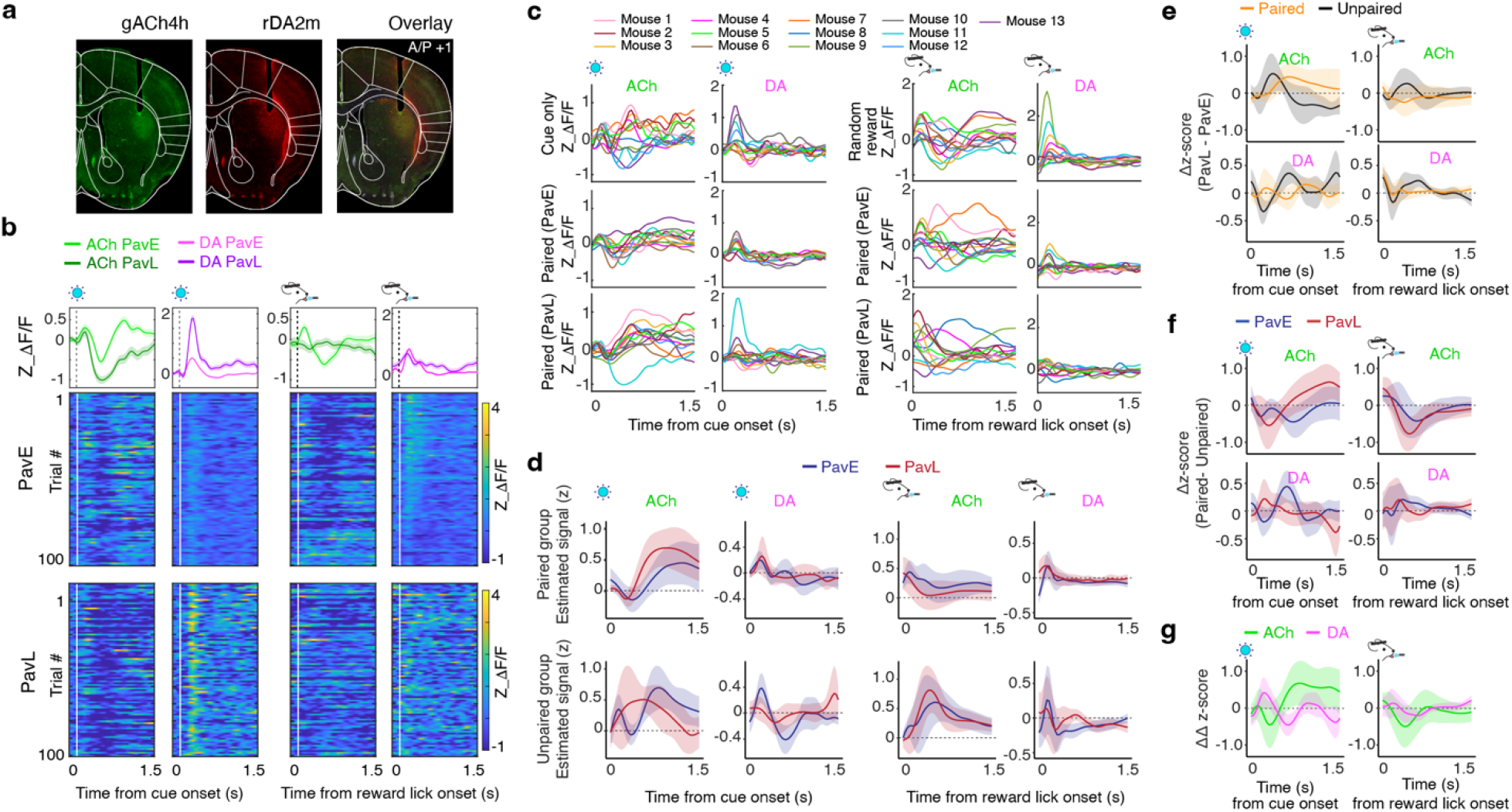
Associative learning differentially reshapes the temporal dynamics of striatal acetylcholine and dopamine. a. Verification of sensor expression and optic-fiber placement shown through histological confirmation of gACh4h (green) and rDA2m (red) expression in the anterior dorsolateral striatum. **b**. Trial-by-trial neuromodulator activity, including representative heatmaps and session averages expressed as mean ± SEM for acetylcholine (ACh) in green and dopamine (DA) in magenta aligned to cue and reward lick onset. Traces compare the first (light) and last (dark) sessions with baseline normalization to 0.5 seconds before cue onset. **c**. Population-level dynamics across conditions representing mean ACh and DA traces aligned to cue (left) and reward lick (right) across cue-only, random reward, and Pavlovian training conditions. Colors represent individual mice in the paired group (n = 13). **d**. Group-level response estimation using bootstrapped generalized additive model (GAM) estimates showing temporal changes in ACh and DA for early (pavE) and late (pavL) sessions in paired (top) and unpaired (bottom) groups. **e**. Learning-dependent shifts defined by the contrast between late and early sessions (pavL - pavE) for paired (yellow) and unpaired (black) cohorts. Left panels display cue-aligned contrasts while right panels display reward-aligned contrasts for ACh (upper) and DA (lower). Shaded areas represent 95% confidence intervals, with deviations from the zero-baseline indicating significant learning-linked modulation of signal magnitude. **f**. Contingency-dependent differences defined by the contrast between training groups (Paired - Unpaired) for early (blue) and late (red) sessions. **g**. Interaction of learning and contingency representing difference-of-differences traces (ΔΔ z-score) that summarize the combined effects of training and group for ACh (green) and DA (magenta).

Across animals, session-averaged traces revealed neuromodulatory signaling features with substantial inter-individual variability in both the paired group (Fig. 2c, n = 13) and the unpaired group (Fig. S3, n = 10). During the earliest training stages (Cue-only, Random-reward), ACh signals were dominated by slow, broad fluctuations with little consistent temporal structure across animals. As learning progressed, paired mice gradually developed a pronounced mid-cue ACh enhancement (duration about 1 sec) that was largely weak in unpaired controls. This emerged alongside the canonical early weak peak and subsequent negative deflection characteristic of predictive cue responses^23,36,37^, but represented a distinct, learning-related cholinergic reorganization. In contrast, DA responses were fast and phasic (<500 ms). Paired mice retained relatively stable cue-evoked DA dynamics, whereas unpaired mice showed reduced amplitudes and increased variability in peak timing and shape by late training. To quantify population-level trends underlying these observations, we fit a generalized additive mixed-effects model (GAMM) and identified two robust motifs (Fig. 2d): (1) ACh signals underwent progressive remodeling, transitioning from slow fluctuations to more structured, multiphasic response profiles as learning consolidated; and (2) DA signals were highly contingency-dependent, preserving stable temporal structure only when a reliable cue–reward association was present.

Model-derived contrasts further isolated the drivers of these changes. Within-group learning contrasts (PavL – PavE; Fig. 2e) revealed a strong dissociation between neuromodulators. For DA, the contrast showed that DA stability depended entirely on predictive validity: unpaired mice exhibited a large negative difference, reflecting the collapse of an initial novelty-driven cue response, whereas paired mice showed minimal change across sessions. ACh learning contrasts instead reflected temporal reorganization: in paired mice, cue-period contrasts showed a clear positive difference corresponding to growth of the mid-cue peak. Between-group contrasts (Paired – Unpaired; Fig. 2f) showed that these differences were modest early in training but became pronounced by late training, especially for ACh, where both cue- and reward-aligned signals developed larger early-phasic negative deflections in paired mice than in unpaired controls.

These divergences were further evaluated by the interaction (difference-of-differences, Learning x Group), which revealed distinct, neuromodulator-specific signatures of learning (Fig. 2g). In the cue period, DA exhibited a robust time-varying interaction characterized by an early positive phase followed by a later negative shift, reflecting the selective stabilization of cue-evoked DA release in paired animals. ACh displayed a qualitatively similar but sign-opposed mean trajectory with an early negative dip and a later positive rebound. However, this cholinergic trend was characterized by greater variability, resulting in confidence intervals that largely encompassed the zero baseline. During the reward period, the interaction was more constrained and temporally limited where ACh signals showed no significant divergence between groups, while DA exhibited only transient, marginal interaction epochs. Thus, the interaction analysis identifies cue-evoked DA stabilization as the strongest contingency-dependent signature of learning, accompanied by a more variable reorganization of cholinergic dynamics. Taken together, these results demonstrate that learning recruits DA and ACh in asymmetric and complementary patterns where DA selectively preserves reward-predictive transients under contingency while ACh undergoes broader, multi-phasic restructuring linked to both cue processing and reinforcement outcome.

### Low-dimensional acetylcholine–dopamine motifs decode trial-wise learning state

We next asked whether trial-by-trial learning state is reflected in the coordinated temporal structure of ACh and DA dynamics, beyond what can be captured by independent scalar features such as peak amplitude or response kinetics. We reasoned that if associative learning reorganizes the temporal relationship between these neuromodulators, the resulting trial-by-trial covariance would more faithfully represent the learned behavioral state than scalar features alone. Supporting this view, low-dimensional representations of population covariance have been shown to capture latent behavioral and cognitive variables with greater fidelity and cross-subject generalizability than scalar measures ^27,29,38,39^. We therefore constructed representations of ACh and DA covariance based on their dominant modes of trial-wise temporal dynamics, termed “motifs,” and tested their capacity to reliably distinguish learned (tcL) from unlearned (tcNL) trials.

To extract these temporally structured response motifs, we performed principal component analysis (PCA) on balanced bootstrapped resamples of cue- and reward-aligned ACh and DA traces across all paired trials (*N* = 16,816; trial counts matched between learned and unlearned states), followed by k-means clustering of the leading principal components (PCs-k; Fig. 3a). ACh signals exhibited greater low-dimensional structure than DA signals, accounting for significantly more variance within the first two PCs (Wilcoxon rank-sum test, *p* < 0.001; Fig. 3b). In the dimension-reduced motif space, reconstructed cue-aligned responses revealed clear divergence between learned and unlearned trials for both ACh and DA, with the separation being more pronounced for ACh (Fig. 3c). By contrast, reconstructed reward-aligned responses showed negligible divergence between learning conditions for either neuromodulator.

**Figure 3.**
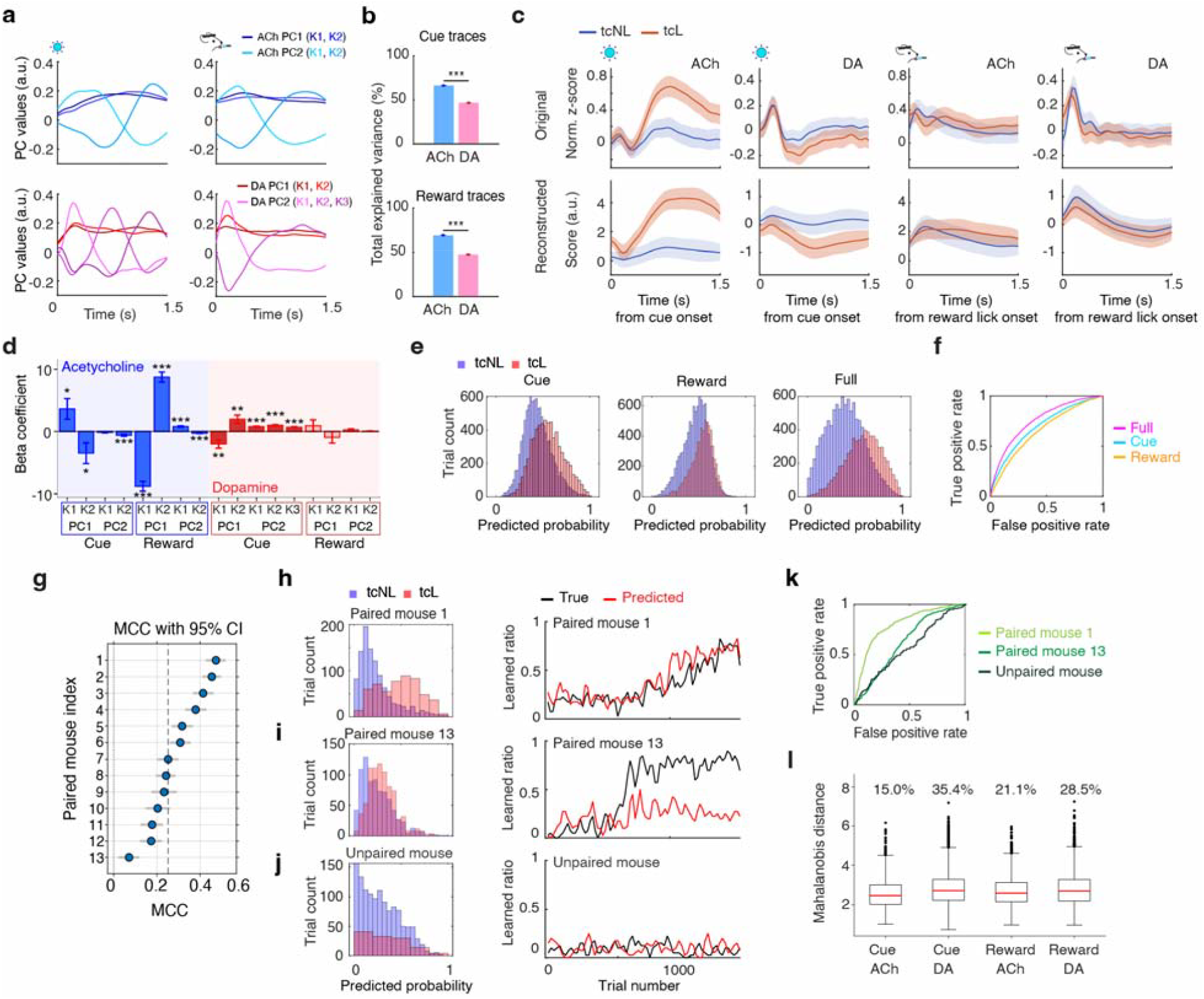
Low-dimensional motif decoding reveals complementary roles of acetylcholine in state geometry and dopamine in learned-state classification. a. Principal component (PC) derived motifs showing the first principal component (PC1) in dark and the second principal component (PC2) in light for acetylcholine (ACh) in the upper panels and dopamine (DA) in the lower panels. Blue and red traces represent signal-specific motifs during the cue period on the left and the reward period on the right where lines represent k-means centroids. **b**. Cumulative variance explained representing the total explained variance for ACh (blue) and DA (pink) across cue-related traces in the upper panel and reward-related traces in the lower panel. **c**. Motif-based signal reconstruction including raw traces (top row) and motif-reconstructed traces (bottom row) for learned trials (tcL) in orange and unlearned trials (tcNL) in blue with shaded regions representing the 95% confidence intervals. The left half of the panel displays cue-aligned data and the right half displays reward-aligned data where the first column contains ACh and the second column contains DA traces. **d**. Logistic-decoder feature of the full decoder based on paired group weights expressed as beta coefficients (β ± standard error) for ACh (blue) and DA (red) on the right in red. Values are shown for cue and reward motifs predicting learned versus unlearned trials where darker bars indicate significantly contributing features. (* *p* < 0.05, ** *p* < 0.01, *** *p* < 0.001). **e**. Decoder performance showing probability histograms for unlearned (tcNL) and learned (tcL) trials across cue, reward, and full-model decoding. **f**. Receiver operating characteristic (ROC) analysis including overlaid ROC curves for the cue (blue), reward (orange), and full (magenta) decoder models. **g**. Leave-one-mouse-out (LOMO) generalization demonstrating decoder performance expressed as the Matthews correlation coefficient (MCC) with 95% confidence intervals (CI) across individual paired mice. **h**. High-generalization example including a probability histogram (left) and trial-by-trial prediction performance compared to behavioral encoding (right) for the highest MCC mouse. **i**. Low-generalization example showing corresponding performance plots for the lowest MCC mouse. **j**. Cross-group validation representing the decoder trained on paired mice applied to an unpaired mouse demonstrating predictions skewed toward the unlearned state. **k**. Cross-group ROC curves comparing individual paired mice and an unpaired control. **l**. Mahalanobis distance between unpaired trials and the paired-group learned-state distribution, including box plots showing the distance-based classification of motifs to confirm motif-specific signal contributions.

To evaluate the functional relevance of these low-dimensional signatures, we trained a logistic regression decoder incorporating all four motif classes (cue- and reward-aligned ACh/DA) in paired mice. At the population level, ACh-related motifs exhibited the largest standardized regression coefficients, indicating consistent cholinergic reorganization across paired animals (Fig. 3d). However, leave-one-mouse-out (LOMO) analyses revealed a dissociation in neuromodulator contributions to decoding performance. Whereas ACh motif contributions remained stable across held-out folds, variability in decoding accuracy was more strongly associated with DA cue-aligned motif contributions (Fig. S4). These findings suggest complementary but distinct roles in learned-state representation: ACh motifs captured broad population-level state transitions, whereas DA motifs contributed more selectively to individual-level discrimination of the learned state.

Decoder performance, assessed by predicted probability and receiver operating characteristic (ROC) analysis, significantly exceeded chance levels when classifying learned versus unlearned trials (AUC = 0.76 ± 0.0001, n = 1,000 bootstrap iterations; Fig. 3e). Notably, the full decoder, integrating both cue- and reward-period motifs from both ACh and DA signals, consistently outperformed single-period decoders, showing a significant improvement in accuracy over models using only the cue period (ΔAUC = 0.059; *p* < 0.001; n = 1,000 iterations) or only the reward period (ΔAUC = 0.101; *p* < 0.001; n = 1,000 iterations). This suggests that neuromodulator dynamics across both task epochs provide non-redundant information about learning state.

The motif-based representation also generalized across subjects. Under a fixed false-positive rate criterion (FPR ≤ 0.30), the median Matthews correlation coefficient (MCC) was 0.25 (interquartile range 0.20–0.39), with all held-out mice exceeding chance levels (Fig. 3g). Despite this overall generalization, individual folds showed substantial performance variability (e.g., MCC = 0.474 in Mouse 1 versus 0.071 in Mouse 13; Fig. 3h,i), consistent with the DA motif dissociation described above (Fig. S4).

We next tested whether the learned-state representation derived from paired mice generalized to the unpaired group. Applying the paired-trained decoder to trials from unpaired mice, without refitting, consistently yielded low learned-state probabilities (Fig. 3j). ROC analysis confirmed that, while even the weakest paired-mouse fold maintained a clearly defined AUC, decoding performance on unpaired trials collapsed toward chance levels (Fig. 3k), indicating that unpaired activity falls outside the associative structure learned from paired conditioning. Furthermore, including unpaired trials in decoder training did not substantially alter model performance, as motifs derived from the full dataset preserved the overall structure and decoding accuracy of the paired-only model (Fig. S5).

To identify which features accounted for the deviation, we quantified the contribution of each motif to the Mahalanobis distance between unpaired trials and the paired-group learned-state distribution (Fig. 3l). Cue-period motifs drove the primary divergence, with the DA cue motif contributing most (35.4% of the total distance) and the ACh cue motif least (15.0%), while reward-period motifs contributed comparably across neuromodulators (DA: 28.5%; ACh: 21.1%).

Collectively, these findings demonstrate that the coordinated low-dimensional temporal structure of joint DA–ACh dynamics robustly encodes trial-by-trial learning state. ACh and DA motifs contributed complementarily but asymmetrically: ACh motifs provided a stable, population-wide scaffold of the learned state, whereas DA motifs contributed more selectively to individual-level discrimination and were the primary feature distinguishing paired from unpaired animals. Critically, the decoder generalized across individuals in the paired group yet classified unpaired trials as “unlearned” despite comparable behavioral performance, revealing a fundamental dissociation between sensory exposure and associative contingency. Whether these two neuromodulator systems also interact directionally on a moment-to-moment timescale is addressed through Granger causality analysis in the following section.

### Learning restructures spontaneous licking and selectively enhances burst-related cholinergic encoding

Beyond cue and reward epochs, paired conditioning reorganized waiting-period spontaneous licking by altering its burst microstructure. Lick rasters revealed a progressive shift from diffuse, sporadic licking to structured, prolonged bouts in paired mice (Fig. 4a top), a transition absent in unpaired controls (Fig. 4a bottom). To quantify this reorganization, we defined lick bursts as consecutive licks with inter-lick intervals < 0.25 seconds (Fig. 4b). Across all animals, spontaneous licking was dominated by brief events with less than 4 licks per burst, providing a data-driven criterion to classify licking into short (≤ 4 licks) and long (> 4 licks) bouts. Although short bursts predominated in both groups, the overall distribution of burst sizes differed significantly between paired and unpaired mice (Wilcoxon rank-sum test, *p* < 0.001; Fig. 4c). A binomial GLMM examining the proportion of short bursts across all 15 training sessions revealed a significantly steeper reduction in the paired group (n = 22) compared with the unpaired group (n = 13; Fig. 4d and e). Post hoc analysis of session-resolved trajectories confirmed a robust effect of training over time in both groups (*F*(1,458) > 357, *p* < 0.001). Crucially, the magnitude of this change was condition-specific, evidenced by a significant Group x Session interaction (*p* < 0.001). While the unpaired group exhibited only a negligible shift in burst composition, the paired group showed a pronounced reorganization of licking behavior (paired: *F* ≈ 24,202; unpaired: *F* ≈ 357). These results indicate that associative learning drives a structural transformation of spontaneous motor output that exceeds the effects of repeated task exposure.

**Figure 4.**
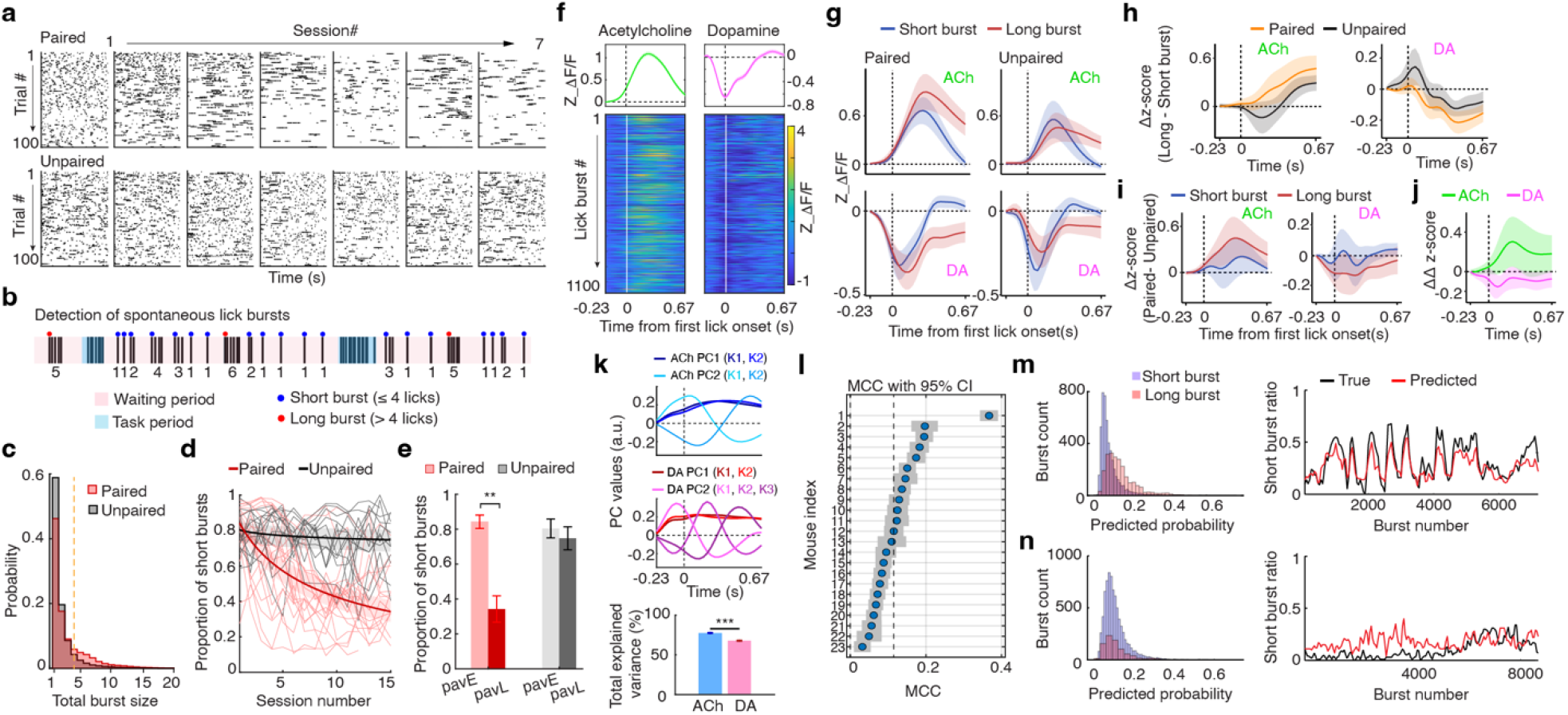
Striatal acetylcholine and dopamine differentially encode spontaneous lick burst size and its refinement with learning. a. Progression of spontaneous licking across sessions representing lick rasters from the waiting period 15 seconds before cue onset for a paired mouse in the top panel and an unpaired mouse in the bottom panel. **b**. Schematics of lick burst classification where spontaneous licks were grouped into bursts if consecutive licks occurred less than 0.25 seconds apart. Bursts were categorized as short for four or fewer licks (blue) and long for more than four licks (red). **c**. Probability distribution of burst sizes demonstrating the probability of occurrence for each burst size in paired mice (red) and unpaired mice (black). **d**. Evolution of burst microstructure showing the longitudinal change in the proportion of short bursts modeled using a binomial generalized linear mixed model (GLMM). Individual thin lines represent data from individual mice, while thick lines represent the group-level fixed effects for paired and unpaired cohorts. **e**. Summary of burst reorganization representing the proportion of short bursts in early (pavE) sessions in light color versus late (pavL) sessions in dark color. Values are expressed as the mean ± SEM for n = 20 paired mice in red and n = 13 unpaired mice in black. **f**. Representative neuromodulator activity at burst onset including mean traces showing acetylcholine (ACh; green) and dopamine (DA; magenta) activity aligned to the first lick of a spontaneous burst at 0 marked by vertical lines in the top panel and a session heatmap in the bottom panel. **g**. Group-level response estimation using bootstrapped generalized additive model (GAM) estimates for ACh (top row) and DA (bottom row). Results are stratified by short bursts (blue) and long bursts (red) for paired mice on the left and unpaired mice on the right. **h**. Burst-size contrasts defined by the difference in signal magnitude between long and short bursts (Long - Short) for ACh in the left panel and DA in the right panel. Contrasts are shown for the paired group in orange and the unpaired group in black where shaded areas represent 95% confidence intervals (CI). Deviations from the zero baseline indicate significant signal modulation by burst duration. **i**. Contingency contrasts representing the difference in signal magnitude between training groups (Paired - Unpaired) for ACh on the left and DA on the right. Contrasts are shown for short bursts (blue) and long bursts (red), with shaded regions indicating 95% CI. Separation from the zero baseline denotes a significant effect of the learning contingency on lick-related signaling. **j**. Interaction of burst size and contingency representing difference-of-differences traces (ΔΔ z-score) that summarize the combined effects of learning and burst size for ACh in green and DA in magenta. Shaded areas represent 95% confidence intervals where periods excluding zero identify significant signatures of behavioral refinement. **k**. Low-dimensional motifs of spontaneous licking including principal component (PC) derived motifs for ACh (blue) and (DA) in red in the top panels and the total cumulative variance explained for each signal in the bottom panels. Motifs represent the first PC1 in a darker color and the second PC2 in a lighter color extracted across iterations. **l**. Leave-one-mouse-out (LOMO) generalization demonstrating decoder performance for burst-size classification across individual mice expressed as the Matthews correlation coefficient (MCC) with 95% CI. **m**. High-generalization example including a probability histogram on the left and burst-by-burst prediction performance compared to true behavioral labels on the right for the highest MCC mouse. **n**. Low-generalization example showing corresponding performance plots for the lowest MCC mouse.

Neuromodulator dynamics at burst initiation exhibited a conserved polarity. Aligning ACh and DA signals to the first lick of each waiting-period burst revealed a rapid DA suppression accompanied by a concomitant ACh elevation (Fig. 4f). To quantify this temporal structure without imposing a priori response window, we fit a GAMM to first-lick-aligned traces from all animals, stratified by burst size and training contingency (Fig. 4g). Model-derived estimates confirmed a robust dependence on burst size, with distinct encoding profiles for each neuromodulator.

Within-group burst-size contrast (Long – Short; Fig. 4h), quantified using varying-intercept mixed-effects bootstrapping (n = 1,000), revealed that long bursts were associated with significantly larger ACh elevations than short bursts. In paired animals, this difference manifested as a pronounced positive deflection occurring throughout the post-lick period. In contrast, DA signals exhibited a consistent negative deflection during long bursts, indicating deeper suppression relative to short bursts in both groups across the duration of the response.

Between-group training contrast (Paired – Unpaired; n = 1,000; Fig. 4i) revealed a clear neuromodulator-specific dissociation where DA responses showed no significant group difference, suggesting a conserved burst-related signal independent of learning (Fig. 4i right). In contrast, ACh signaling was strongly modulated by training history as paired animals exhibited sustained elevations relative to unpaired controls (Fig. 4i left). These differences were prominent during the later phases of the lick-locked traces for both short and long bursts. Finally, to determine whether training altered the neural encoding of burst size, we quantified the difference-of-difference interaction term. This analysis revealed a significant positive interaction for ACh within the post-lick window, which indicates that associative learning selectively enhances the gain of ACh burst-size encoding (Fig. 4j). In contrast, DA showed no significant interaction effect, reinforcing the interpretation that DA scaling reflects a conserved, training-independent signature of lick initiation.

We next tested whether burst size is decoded in the low-dimensional covariance of neuromodulator dynamics. We extracted PCA-derived motifs from bootstrapped, and first-lick– aligned traces revealed that ACh motifs explained more variance than DA motifs (variance explained by PC1-2: ACh 77.83 ± 0.26% (CI) vs. DA 68.12 ± 0.31% (CI); *p* < 0.0001; Fig. 4k). However, motif-based decoding of burst size exhibited limited cross-animal generalizability, with modest overall performance (median MCC = 0.12, IQR = 0.07 – 0.15, n = 23; Fig. 4l). The best-performing fold (MCC = 0.367; Fig. 4m) exhibited separability in predicted burst-size probabilities and a close correspondence between decoded states and the temporal evolution of burst structure. In contrast, the weakest-performing fold (MCC = 0.032; Fig. 4n) showed substantial overlap between predicted probability distributions and poorly tracked transitions in burst structure.

Together, these results demonstrate that while spontaneous lick initiation triggers a conserved, training-independent DA suppression, ACh signaling undergoes a fundamental reorganization that selectively amplifies information about motor vigor during associative learning. This divergence between the two neuromodulators suggests that the cholinergic system provides a flexible, state-dependent scaffold for motor output whereas DA maintains a consistent signature of action initiation.

### Dopamine exerts a sustained Granger-causal influence on acetylcholine

To determine whether moment-to-moment fluctuations in striatal DA and ACh carry directional predictive structure, we performed bivariate Granger causality (GC) analysis on simultaneously recorded session-long photometry signals (Fig. 5a). We first confirmed that GC estimates did not vary significantly across Pavlovian learning stages or session order. Consequently, data from the first and last training sessions were pooled to maximize statistical power (n = 44 sessions, 22 mice, from paired and unpaired groups). To minimize spurious inference from shared slow drifts, we selected autoregressive lag order using residual whitening (Ljung–Box criterion) and AIC improvement, ensuring removal of residual autocorrelation within 1 s.

**Figure 5.**
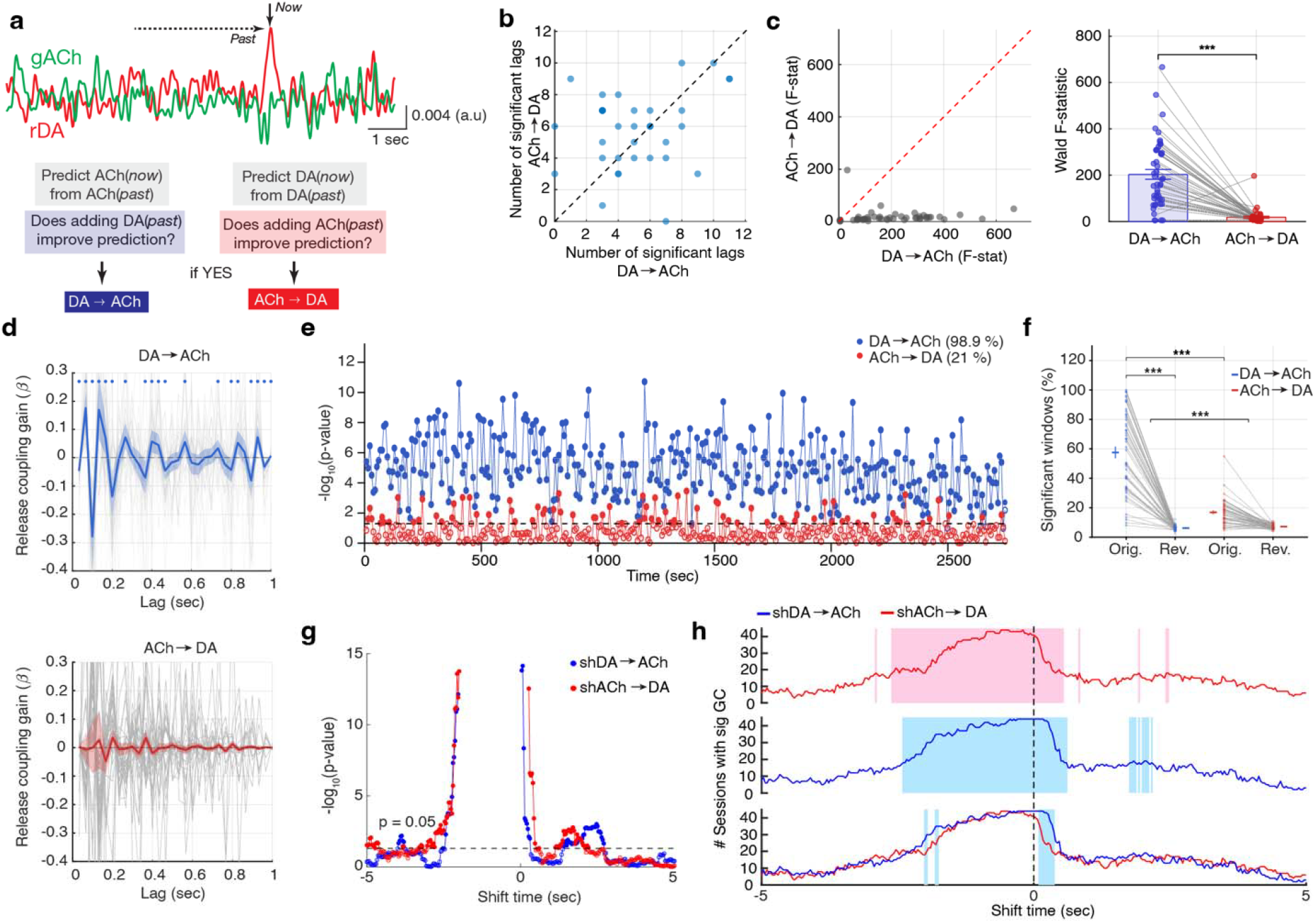
Granger causality analysis reveals an asymmetric temporal influence of dopamine on acetylcholine dynamics. a. Schematic of temporal predictive relations between dopamine (DA) and acetylcholine (ACh) showing 15 seconds of ACh (green) and DA (red) traces. This panel illustrates the concept of Granger causality (GC) within this data, highlighting how current signal values are predicted using both their own history and the past values of the other neuromodulator. **b**. Session-wise comparison of the number of significant cross-lag regression coefficients in the DA→ACh versus ACh→DA directions. Points clustering around the identity line indicate a comparable number of significant lagged influences in each direction. **c**. Scatter plot of F-statistics (left) and paired comparison (right) showing higher Wald F-values for DA → ACh (blue) than ACh → DA (red) across sessions. **d**. Lag-resolved impulse response functions (IRFs) showing structured temporal influence across lags for DA → ACh (upper panel; blue) and ACh → DA (lower panel; red). Dots indicate individual lags that exhibited population-level significance as determined by percentile bootstrapping of the mean regression coefficients. Shaded regions represent 95% bootstrap confidence intervals where the DA → ACh direction shows structured predictive power while ACh → DA intervals consistently overlap with zero. **e**. Example session showing the temporal progression of significant GC windows computed using a 15-second window size and a 7.5-second sliding step. Filled circles indicate *p* < 0.05 relative to the dashed significance threshold, and empty circles represent non-significant windows. The percentage of significant windows per session was used as the primary metric to quantify the temporal stability of these predictive relationships. **f**. Temporal specificity control comparing the percentage of significant windows between original and time-reversed data. The left blue pair represents the DA → ACh comparison while the right red pair represents the ACh → DA comparison. Statistical tests evaluated differences between original data across directions and between original and reversed data within each direction as well as the difference-of-differences between the two directions (*** *p* < 0.001). **g**. Systematic lag-shift analysis of an example session with predictor signals shifted systematically showing significance plotted across lag shifts from -5 to +5 seconds. Samples were converted to time based on a 30 Hz sampling rate for shifted dopamine (shDA) in blue and shifted acetylcholine (shACh) in red. **h**. Directional lag-shift distributions and population significance. *Top and Middle:* Number of sessions with significant GC across lag shifts for shACh → DA (red) and shDA → ACh (blue). Shaded regions indicate lag shifts exceeding the permutation-tested significance threshold. *Bottom:* Directional comparison between the two pathways. Blue shading highlights lag windows where the count of significant sessions for shDA → ACh significantly dominates over shACh → DA (*p* < 0.05, permutation test).

We first compared the temporal extent of directional interactions by quantifying the number of significant cross-lag regression coefficients in each direction. Session-wise comparisons clustered around the identity line (Fig. 5b), indicating a comparable number of detectable lagged influences between DA → ACh and ACh → DA directions. However, the strength of directional influence was strongly asymmetric. Session-wide Wald F-statistics, reflecting overall predictive gain, were consistently larger for DA → ACh than for ACh → DA (*p* < 0.001, paired t-test; Fig. 5c), demonstrating that DA history provides substantially greater forward predictive influence on ongoing ACh dynamics than the reciprocal cholinergic contribution. To resolve the fine-scale temporal organization of coupling, we examined lag-resolved impulse response functions (IRFs). The DA → ACh model exhibited structured, higher-amplitude coefficients, with multiple lags reaching population-level significance via percentile bootstrapping of the mean regression coefficients. Within this causal drive, significant positive coupling gain was concentrated during the early response phase and was followed by a transient period of negative coupling at intermediate lags (Fig. 5d top). In contrast, ACh → DA coefficients were comparatively small and did not to reach significance at any lag as the 95% bootstrap confidence intervals consistently overlapped with zero, reinforcing the unidirectional nature of the causal drive (Fig. 5d bottom). In shuffled control datasets, these directional patterns were abolished (Fig. S6), confirming that the observed predictive relationships depend on the temporal structure of the signals.

We evaluated the temporal stability of this asymmetric hierarchy using sliding-window GC analysis (15-s window, 7.5-s step) across learning stages in both paired and unpaired mice. In a representative session, DA → ACh interactions remained significant across 98.9% of windows, whereas ACh → DA effects were intermittent (21%; Fig. 5e). This disparity was highly consistent across the population; the proportion of significant windows was significantly greater for DA → ACh (60.8 ± 4.0%) than for the reverse (17.4 ± 1.3%; Wilcoxon signed-rank test, *z* = 5.60, *p* < 0.0001; Fig. 5f). Importantly, time-reversal controls abolished significant coupling in both directions (DA original vs. reversed: *p* < 0.0001; ACh original vs. reversed: *p* < 0.0001), confirming that the predominant dopaminergic influence arises from predictive temporal structures rather than shared signal properties. Furthermore, a bootstrap-derived interaction test (10,000 iterations) confirmed that the reduction in significant engagement following time-reversal was significantly larger for the DA → ACh drive than for the reciprocal feedback (mean DA drop: 54.5% vs. mean ACh drop: 10.2%; *p*_*diff*_ < 0.0001; Fig. S6c), underscoring the temporal specificity of the dopaminergic causal drive. Because the 15-s analysis window is substantially shorter than individual trial duration, GC windows were distributed across both cue/reward epochs and off-task waiting periods, indicating that the DA→ACh asymmetry reflects a persistent cross-state coupling intrinsic to the circuit dynamics.

To further characterize the temporal specificity of this interaction, we performed a systematic lag-shift analysis by offsetting the predictor signal relative to the target across ±10 s (30 Hz sampling rate). In both directions, predictive strength peaked near zero shift and declined asymmetrically, with significance persisting longer at negative lags (Fig. 5g). Population-level permutation testing confirmed that significant sessions were robustly concentrated near zero shift, spanning approximately −3 s to 1 s (Fig. 5h top and middle). Direct comparison of significant session counts, however, revealed a clear directional divergence. DA → ACh exhibited a significantly greater predictive influence across zero-centered lag shifts than the ACh → DA with the strongest difference in the shift range of 0 – 500 ms (Fig. 5h bottom).

Collectively, these findings support a robust and stable directional hierarchy in which DA fluctuations serve as a primary predictive scaffold for striatal ACh dynamics, identifying DA as a dominant driver of moment-to-moment coordination in the striatal neuromodulatory landscape.

## Discussion

We investigated the temporal dynamics of DA and ACh in the aDLS during associative learning using longitudinal dual-color fiber photometry combined with a multi-scale analytical framework spanning session-averaged, trial-resolved, and moment-to-moment timescales. Across training, animals developed anticipatory cue responses and suppression of spontaneous licking selectively in the paired condition, marking a shift toward a learned behavioral state. This transition was accompanied by distinct neuromodulator dynamics: DA maintained fast, contingency-dependent cue transients, whereas ACh underwent broader temporal restructuring, developing a pronounced mid-cue enhancement. Beyond average responses, low-dimensional motif analysis revealed that learning state was embedded in the trial-by-trial temporal structure of joint DA–ACh dynamics. A decoder constructed from cue- and reward-related neuromodulator motifs reliably identified trial-wise learned states and distinguished paired associative learning from unpaired sensory exposure. Learning also reshaped waiting-period spontaneous licking into long-burst patterns and selectively enhanced cholinergic encoding of lick-burst vigor, whereas DA maintained a training-independent signature of action initiation. Granger causality analysis further revealed a short-latency asymmetric predictive influence from DA to ACh. This directionality persisted across learning stages in both paired and unpaired mice, indicating a stable temporal hierarchy between DA and ACh. Together, our findings identify DA and ACh not as independent striatal signals, but as a coordinated neuromodulatory system in which DA provides a leading temporal scaffold and ACh carries broader state- and action-related reorganization during learning.

Rodent licking behavior exhibits structured burst patterns and training can lead to the emergence of anticipatory licking prior to reward delivery^40^. Moreover, reinforcement-learning frameworks show that behavioral policies adapt across training^41^. Spontaneous licking outside task-relevant epochs declined with training, indicating that animals learned to suppress indiscriminate licking and prioritize cue-driven responses. This suppression paralleled the emergence of anticipatory cue-evoked licking, which became the dominant behavioral mode under pairing conditions^42^. Within this trajectory, we have observed that paired mice selectively pruned short bursts while retaining long bursts during waiting period, which may represent provisional “practice units” observed during refinement of learned behavioral motifs^42^. Consistent with reports that mice preferentially re-express behavioral syllables previously associated with reward, these spontaneous long bursts likely reflect transient rehearsal forms expressed during the learning process. These bursts eventually diminished with extended training, suggesting they constitute an intermediate stage in the refinement of anticipatory behavior.

The behavioral refinement was accompanied by distinct neuromodulator dynamics. Using conventional scalar analyses, we captured lick-aligned DA and ACh signaling through amplitude and waveform changes, with a transient negative deflection of DA started before lick initiation accompanied by an anti-correlated phasic increase of ACh, which is consistent with reports that movement-related variance dominates dorsal striatal signals^38,43^. Yet decoding the size of spontaneous licking burst from low-dimensional motifs of these neuromodulators was modest. By contrast, cue- and reward-related differences appeared subtle in scalar metrics —likely due to cross-animal variability and multiple contributing factors—yet supported robust decoding of learned state from a small set of low-dimensional components. This dissociation suggests that burst size is primarily encoded through response magnitude and shape, whereas learned state is embedded in coordinated, low-rank covariance structure of DA and ACh. This is consistent with evidence that structured neural population dynamics capture latent cognitive variables beyond average response levels^27,44^.

Dual-color photometry studies in the dorsal striatum have revealed coordinated DA and ACh fluctuations during reward and movement epochs^14,26^, and motor learning has been shown to shift DA responses from reward consumption toward the cue period^45^. The accompanying transient dip in ACh during reward-predictive cues is only partially dependent on D2-receptor signaling^17^. Beyond this previously reported phasic suppression, our session-averaged ACh responses revealed an additional temporal feature: a slow, broad secondary peak during the cue period, which is clearly distinct from both the initial weak ACh peak and the faster DA transient^45,46^. An interaction contrast (Session x Group) further identified a mid-cue positive rebound in ACh arising selectively in paired animals, which becomes more significant when comparing ACh signaling in learned vs. unlearned trials. This rebound is consistent with reports linking sustained cholinergic activity to effort-related signaling in the ventral striatum ^47^ and with multiphasic ACh responses observed in the dorsal striatum following extinction^36^, suggesting that it may reflect a learning-dependent reconfiguration of cholinergic gain during cue-reward association.

Trial-wise low-dimension analysis further extended these findings by showing that learned-state information is jointly represented across both neuromodulators, with a stronger and more consistent contribution from DA. Rather than uniform gain changes, learning appears to selectively reorganize a coordinated low-dimensional subspace spanning DA and ACh, most prominently during cue-related processing, which carries learned-state information, while movement-related variance is more closely linked to the ongoing temporal structure of the motor output. This reorganization may reflect or contribute to synaptic plasticity at corticostriatal inputs onto SPNs^48,49^, where DA and ACh exert complex and interacting influences^50,51^. Such a view explains cross-animal variability without invoking global signal changes and links behavioral refinement to ensemble-level remodeling. The observation that the cholinergic interaction exhibits a sign-opposed mean profile relative to DA but with greater variability suggests that while the underlying trend is conserved, the cholinergic system undergoes a more heterogeneous reorganization than the dopaminergic system. We therefore interpret the stimulus-locked low-rank consolidation as a systems-level signature of learning, which is compatible with SPN plasticity but not reducible to scalar modulation in either channel alone. More broadly, the complexity and non-uniformity of this reorganization may be especially well suited to the aDLS’s role in habitual learning, where behaviors are shaped gradually through extended trial-and-error^52,53^.

Decoder performance generalized across most paired animals, though in one case the separation of learned from unlearned trials was relatively poor compared to other subjects while remaining above chance levels. Such differential generalizability suggests that trial-level motif representations reflect variable and subject-specific reorganization of the ACh/DA dynamics even within a common behavioral paradigm. One explanation is that our framework is sensitive to latent biological heterogeneity, such as differences in striosome–matrix organization, which is known to influence DA and ACh release dynamics^54-57^. Alternatively, decoding structure may diverge with distinct learning trajectories, with rapid learners compressing or bypassing intermediate motif states^42^. In this view, lower decoder performance does not indicate noise but a true shift in underlying computation. Thus, motif-based decoding provides a potential window into both subregional specialization and individual variability in neuromodulatory engagement across learning.

By applying physiologically constrained Granger causality analysis during associative learning— consistent with methodological guidance from Stokes and Purdon^30^— we uncovered a temporal asymmetry: DA fluctuations reliably precede and predict subsequent ACh activity, whereas the reverse relationship is substantially weaker. This directional asymmetry aligns with established circuit motifs, including temporally structured DA–ACh interactions observed in slice optogenetic studies^20,58^ and the spatially organized properties of DA terminals^59-61^. Previous work by Krok et al^23^ demonstrated that spontaneous striatal DA and ACh fluctuations maintain a consistent temporal relationship yet are not coordinated through direct local intrastriatal interactions. Our results do not contradict this framework at the level of necessity but reveal an additional dimension of their relationship. Although DA and ACh fluctuations might not require direct local interactions to co-occur, their moment-to-moment dynamics are nonetheless temporally asymmetric. This directional asymmetry persisted across full recording sessions in both paired and unpaired animals and was reproduced during off-task waiting periods free of explicit cue or reward events, indicating that it reflects an intrinsic feature of circuit dynamics rather than a transient structure imposed by task engagement. Together, these results suggest that while neither signal is strictly required for the occurrence of the other, DA provides a predictive temporal scaffold that precedes ACh dynamics across behavioral states — an organizational feature not detectable by loss-of-function approaches but evident in fine-grained temporal dependencies.

In summary, our study demonstrates that striatal DA and ACh show distinct yet complementary dynamics during associative learning. Behavioral refinement was paralleled by plasticity in cue- and reward-evoked signals, but learning state was more accurately captured in low-dimensional covariance motifs than in average response amplitudes. Motif-based decoding revealed that DA contributed the most consistent and learning-sensitive temporal structure, whereas ACh exhibited coordinated but more variable dynamics across trials. Granger causality further uncovered a stable asymmetry in which DA exerted a strong predictive influence on ACh - a temporal scaffold that precedes and predicts ACh dynamics across behavioral states. By combining longitudinal dual-color photometry with advanced decoding and causality analysis, this work establishes a framework for quantifying how multiple neuromodulatory signals jointly encode behavioral state. More broadly, it demonstrates how low-dimensional motifs and temporal asymmetries can reveal circuit principles not evident from conventional averages or ablation studies. Such an integrative view may help explain how DA and ACh contribute in concert to striatal computation and guide future studies on how their interactions are reorganized in disease^62,63^.

## Methods

### Animals

Experimental mice were Cre-negative male and female offspring bred in-house from A2a-Cre or D1R-Cre males (Jackson Laboratory) crossed with C57BL/6J wild-type females. Genotyping was performed by Transnetyx, and only Cre-negative mice were used. Animals were 11–13 weeks old at surgery. Animals were housed on a 12 h light/dark cycle with ad libitum food and water unless noted. All procedures adhered to institutional and national guidelines and were approved by the Institutional Animal Care and Use Committee at the University of Texas Southwestern Medical Center.

### Mouse preparation

Anesthesia was induced with 3–4% isoflurane and maintained at 1–2%. Mice were placed in a stereotaxic frame on a heating pad (37 °C). Slow-release buprenorphine was given subcutaneously for analgesia. After hair removal and scalp incision, the skull was cleaned and etched with 3% hydrogen peroxide. Burr holes were drilled above the anterior dorsolateral striatum (aDLS) relative to bregma (AP +1.0 mm, ML +2.1 mm, DV +3.3 mm). A mixture of GPCR-based viral sensors for acetylcholine (AAV9-hsyn-gACh4h) and dopamine (AAV9-hsyn-rDA2m) was injected into the right hemisphere (250 nL at 2 nL/min; titer ∼2 × 10□ copies each). The pipette was left in place for 5 min to minimize reflux. An optic fiber was placed 0.5 mm above the injection site, and the craniotomy was sealed with Kwik-Sil or tissue adhesive. Metabond was applied to the skull and fiber, and a lightweight 3D-printed headbar was affixed with dental cement. After recovery from anaesthesia, mice were returned to their home cage on a heating pad. Post-operative care included daily monitoring and carprofen injections for three days.

### Behavioral learning paradigm

Experimental cohorts consisted of both male and female mice which were randomly assigned to either the paired or unpaired training conditions. To minimize potential sibling effects, subjects were selected from distinct litters so that minimal individuals from the same pup cohort were represented within any single experimental group. A total of 20 paired and 13 unpaired mice completed all behavioral training sessions. Our analysis of learning progress and burst proportion changes utilized data from this full cohort (*N* = 33). For all photometry-related analyses, we included only those subjects with histologically verified fiber placement and sensor expression in the targeted anterior dorsolateral striatum which consisted of 13 paired and 10 unpaired mice.

Three weeks after viral expression and fiber implantation, signal quality was tested during a 1-h head-fixed session. If signals were suboptimal, mice were held for an additional week; expression quality was reassessed weekly up to six weeks, after which unsuccessful cases were excluded and verified histologically.

Once acceptable signals were obtained, mice were placed on a water regulation schedule. Hydrogel (99% water) was provided at 4% of body weight daily, gradually reduced over one week to establish motivation. Before reward exposure, mice completed a “cue-only” session of 40 LED cue presentations to familiarize them with the visual stimulus.

Mice then underwent reward habituation, receiving 4 μL of 4% sucrose every 30 s. Initially the spout was positioned near the mouth, then gradually moved to a fixed location requiring tongue protrusion. After consistent reward retrieval, mice completed a “free reward” session, where reward was delivered if they licked within 4 s of trial onset. With variable inter-trial intervals (8– 12 s), most mice obtained >200 rewards within 1 h. Those below this threshold repeated the session until reaching criterion.

Next, mice completed 1–2 “random reward” sessions (ITI = 20–30 s), receiving up to 100 rewards/session without cues. From this stage onward, animals were randomly assigned to paired or unpaired groups, balanced by sex and cohort.

Paired mice underwent 16 training sessions. The first included 40 unpaired followed by 60 paired trials. Paired trials consisted of a 1.5-s LED cue immediately followed by sucrose delivery (4 μL, 4%). Unpaired mice received 15 sessions of unpaired trials, where cues and rewards were decoupled, with reward delivered randomly during the ITI (22–32 s). Fiber photometry was recorded during all sessions except sucrose habituation.

Motivation was dynamically adjusted. If lick rate fell below 35/min, new trials were paused until 12 spontaneous licks were detected, after which trials resumed. Sessions ended at whichever came first: 100 rewards, 120 trials, or 1 h duration.

### Clustering cue-elicited licking patterns to index Pavlovian learning stages

During Pavlovian training, licking activity during the pre-cue and cue periods reflected distinct behavioral states. To quantify these patterns, cumulative lick counts and lick intervals were computed within a −1.5 to +1.5 s window around cue onset. A second-order polynomial was fit to these data, with cumulative lick interval as the independent variable and cumulative lick count as the dependent variable, capturing both first- and second-order features of lick timing.

Trials were categorized into four clusters based on the fitted curves. Cluster 0 included trials in which no licking occurred during the analyzed period. Cluster 1 consisted of trials with some pre-cue licking but weak cue-evoked responses. Cluster 2 showed minimal pre-cue licking but strong, early cue-evoked licking. Cluster 3 did not exhibit a consistent temporal structure; licks occurred sporadically without clear pre-cue or cue-locked responses.

For subsequent analyses, Cluster 0 and Cluster 3 were grouped as unlearned trials (tcNL), whereas Cluster 1 and Cluster 2 were considered learned trials (tcL), reflecting the emergence of anticipatory licking associated with cue–reward learning.

### Spontaneous lick adaptation during Pavlovian learning

As cue-evoked anticipatory licking emerged during Pavlovian training, spontaneous licking outside task epochs occurred. During this adaptation, many mice emitted bursts of licks during waiting periods, prompting a more detailed analysis of burst structure.

We defined spontaneous licking as licks occurring outside cue and reward epochs (from reward delivery to the end of consummatory licking). Licks separated by <250 ms were grouped into a bout. Bouts were categorized as short (≤4 licks) or long (>4 licks). For each session, we quantified spontaneous lick rate, the proportion of short vs. long bursts, and their evolution across training.

### Dual-color fiber photometry

We built a dual-color fiber-photometry system capable of simultaneous recording in up to four independently operated behavioral chambers. Each chamber was controlled by a dedicated Raspberry Pi and Arduino for peripheral devices and TTL synchronization. A fixed optic fiber collected fluorescence signals from head-fixed mice during behavioral tasks.

Each chamber was connected to a 6-port miniCube (iFMC6-G2; Doric Lenses), which combined excitation and emission paths. Excitation light included 405 nm (isosbestic control), 470 nm (active green channel for gACh4h), and 565 nm (active red channel for rDA2m), each delivered through separate ports. Importantly, the two sensors exhibit comparable kinetics (rDA2m: τ_on = 50 ms, τ_off = 2.24 s; gACh4h: τ_on = 70 ms, τ_off = 2.10 s; data courtesy of Dr. Yulong Li), supporting direct comparison of temporal profiles across the two channels and analyses of DA–ACh coordination during learning. Light traveled via a sampling cable into the brain and emission signals returned through the same path. The miniCube filtered emission into green and red channels, with built-in photodetectors converting fluorescence to voltage outputs.

Each LED was driven by an independent LED driver (Thorlabs). One complete set included three drivers (405, 470, 565 nm), and four identical sets supported four chambers. Timing of excitation was controlled by TTL pulses: LEDs were sequentially triggered for 1.5 ms in the order 405–470–565 nm, with 11 ms spacing, repeating continuously. This resulted in an effective sampling rate of 30 Hz per wavelength.

Analog voltage signals from the photodetectors, together with excitation TTL pulses and trial start markers from four chambers, were digitized using a LabJack T7-Pro DAQ (CB37 terminal board). Eight analog channels recorded green and red emission signals from each chamber. A single digital channel recorded synchronization TTL inputs from seven ports, each assigned a unique 2^n^-weighted value, allowing simultaneous detection of multiple event sources. Port identity was extracted by binary decoding of the recorded digital values. All signals were acquired at 10 kHz, then downsampled and parsed offline into per-chamber datasets aligned with trial events.

Post-processing extracted peak values from each excitation pulse (405, 470, 565 nm), generating three raw traces per emission channel. This yielded six datasets per session (three for green, three for red). Traces were low-pass filtered at 4 Hz to remove high-frequency noise, then z-scored. For green fluorescent protein (GFP) based sensors, ΔF/F was computed by referencing each 470 nm sample to its reference 405 nm sample: (X470 − X405) / X405. ΔF/F traces were subsequently z-normalized.

To align neural signals with behavioral events, timestamps were synchronized between the photometry and behavioral control systems. Trial start TTLs were used to calibrate timing offsets, and each behavioral event was matched to the nearest photometry timestamp. Event-aligned traces were extracted for cue onset/offset, first lick after reward, and spontaneous licking bouts.

For cue- and reward-aligned traces, a 2 s window (−0.5 to +1.5 s relative to the event) was extracted, and baseline correction was applied using the 0.5 s pre-cue period. For lick-aligned traces, a 0.9 s window (−0.21 to +0.69 s around lick onset) was used, with the first extracted value served as baseline.

### Functional Regression Analysis

To characterize the temporal evolution of event signals for DA and ACh during learning, we applied functional regression modeling using the *pffr* function from the refund R package. Trial-wise baseline calibrated traces were modeled for three event types, including cue-aligned (45 timepoints), reward-aligned (45 timepoints), and lick-aligned (28 timepoints) traces based on the first lick of each burst.

#### Model structure

For cue- and reward-traces, the functional outcome was expressed as:

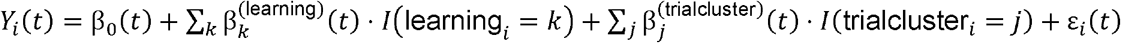

In this equation, *Y*_*i*_ (*t*) represents the trace at time *t* while β_0_ (*t*) is the functional intercept. The terms β^(learning)^ and β^(trialcluster)^ represent smooth functions for learning stage and trial type (learned versus not learned), *I* denotes the indicator function, and ε_*i*_ (*t*) represents residual error.

For lick-aligned traces, we included an additional predictor for burst size.

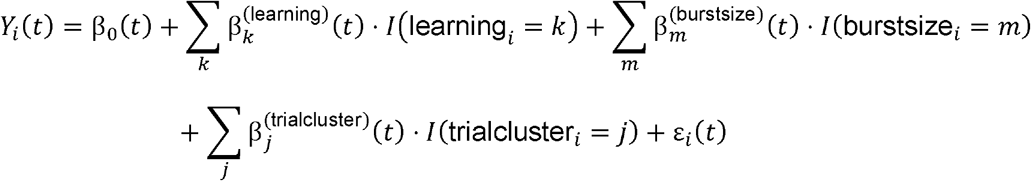

In this model, burst size was coded as a binary factor of short versus long bouts. Penalized B-splines were utilized with a basis dimension of *k*= 25 to ensure sufficient flexibility while preventing overfitting. Models were fit using the bam engine to optimize computational efficiency for large trial-wise datasets.

#### Experimental factoring and condition classifications

For paired mice, cue-aligned signals were modeled across five learning stages including cue-only, unpaired and paired trials within Pavlovian Day 1, early Pavlovian (pavE), and late Pavlovian (pavL). Each trial was further classified as learned or not learned based on the lick cluster analysis. For unpaired mice, models included only three stages consisting of cue-only, pavE, and pavL with no trial classification.

Reward-aligned traces for paired mice were modeled across random reward, Day 1 unpaired and paired trials, pavE, and pavL. Trial classification was defined by licking behavior during the preceding cue rather than the reward period itself. This allowed reward responses to be interpreted in the context of the preceding cue association. Unpaired mice were modeled across random reward, pavE, and pavL only.

Lick-trace modeling included three learning stages consisting of random reward, pavE (licks from the first half of training), and pavL (licks from the last half). Predictors included burst size and the trial cluster based on the preceding cue period. Bursts arising after a learned trial were classified as belonging to a learned cluster to capture how licking dynamics were shaped by both burst structure and learning context.

### Bootstrap procedure and interaction inference

We implemented a bootstrap procedure consisting of 1000 iterations to generate condition-specific mean traces and population-level estimates. Because the functional regression was fit at the individual level, this resampling approach allowed us to account for inter-subject variability when deriving population contrasts. We computed pairwise differences for within-group learning effects and between-group session effects.

The interaction (difference-of-differences) was calculated by subtracting the unpaired learning contrast from the paired learning contrast. We derived 95% confidence intervals at each time point and interpreted periods where the confidence interval excluded the zero baseline as evidence of significant interaction-like effects. This framework enabled the dynamic tracking of neuromodulatory signals while isolating stage-dependent changes and trial-level variability across cue, reward, and lick responses.

### Neural Decoder Construction and Evaluation

We developed a logistic regression decoder using low-dimensional waveform motifs extracted from raw neural traces to test how striatal neuromodulatory dynamics relate to behavior. The pipeline comprised three primary stages including motif extraction, decoder training, and cross-animal validation.

#### Motif extraction

Cue and reward-aligned trial-wise ACh and DA traces as well as spontaneous lick-aligned traces were compiled across mice and sessions. For each signal-event combination, we applied a bootstrapped PCA-k-means procedure. In each of 500 iterations, 80 traces were randomly sampled and balanced between learned and unlearned trials for cue and reward or between short and long bursts for licking. PCA was then performed and the first two principal components (*PC*1 and *PC*2) were retained. K-means clustering with *k* ≤ 3 identified motif patterns within this reduced subspace where the optimal *k* was determined by elbow and silhouette criteria.

Cluster centroids were projected back into the time domain and normalized to yield motif templates. These templates consisted of DA and ACh *PC*1 – *k* and *PC*2 – *k* patterns, which represented the dominant temporal motifs across iterations. Each trial or lick burst was subsequently scored against these templates using normalized dot products. This generated motif similarity scores that served as the primary predictors for downstream decoding analyses.

#### Decoder training

Motif similarity scores were used as predictors in logistic regression models fit using a stepwise generalized linear modeling procedure with a binomial link. For the cue and reward decoders, the response variable was trial class (learned versus unlearned) defined according to cue-period licking. For the lick decoder, the response variable was burst size (short versus long). Both main effects and two-way interactions were considered for inclusion. Predictors were retained if they significantly improved model fit as measured by a reduction in the Akaike information criterion (AIC).

#### Cross-animal validation and performance evaluation

Generalization was tested using a leave-one-mouse-out (LOMO) framework. For each fold, motifs were extracted from all mice except for one held-out subject and models were trained using the corresponding similarity scores. A fixed decision threshold was chosen on the training set by maximizing the Matthews correlation coefficient (MCC) under the constraint of false positive rate (FPR) ≤ 0.30. The same threshold was then applied to the held-out mouse.

Performance was summarized primarily by MCC with additional metrics including recall, precision, balanced accuracy, FPR, receiver operating characteristic area under the curve (ROC-AUC), and precision-recall area under the curve (PR-AUC) for reference. Sliding-window analyses were further utilized to track the temporal evolution of predicted learned trials for cue and reward periods or burst-type predictions for licking behavior.

### Granger Causality Analysis

We assessed temporal interactions between ACh and DA signals using Granger causality (GC) with dynamic regression models. This analysis tests whether the past values of one signal improve the prediction of another beyond the predictive power of its own history.

#### Preprocessing and lag structure selection

Time series were detrended with polynomial regression and mean-centered before analysis. Autocorrelation function (ACF) and cross-correlation function (CCF) plots guided the initial lag selection. The CCF results consistently suggested that lagged DA predicted current ACh which supported the presence of directional temporal interactions.

We utilized a hybrid procedure combining residual whitening and an AIC-based search to determine the optimal lag structure. The self-lag length was first established by Ljung-Box tests to ensure that residuals were free of autocorrelation up to 30 lags which corresponds to approximately 1 second. Cross-lags were then added sequentially and were retained if they improved the model fit as indicated by a Δ*AIC* ≥ 4. This selection process stopped if five consecutive lags failed to improve the fit or reached a maximum of 30 lags. A 20% buffer of self-lags was added to the final model to safeguard mathematical stability.

#### Model fitting and statistical testing

Dynamic regression models were fit by ordinary least squares according to the following equation.

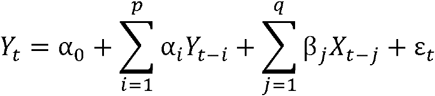

In this model, *Y*_*t*_ represents the response signal such as DA while *X*_*t*-*j*_ represents the predictor cross-lags such as ACh. The terms *p* and *q* denote the self-lag and cross-lag numbers respectively while ε_*t*_ represents the residual error. The full model included both self-lags and cross-lags while the restricted model contained only self-lags. Significance of individual coefficients was assessed using a Bonferroni correction at a threshold of *p* < 0.05.

We tested for predictive influence using a Wald F-test to compare the full versus restricted models. A significant improvement in the model fit indicated a predictive relationship between the signals. All analyses were performed bidirectionally to evaluate both *ACh* → *DA* and *DA* → *ACh* influences.

#### Temporal structure controls and sliding window analysis

To rule out spurious results due to shared rhythms, we performed a time-reversal control. The predictor signal was reversed in time while the response signal remained intact and models were refit with identical lag structures. We interpreted Granger causality effects that were present in the original data but absent under time-reversal as genuine directional predictions.

To examine the temporal stability of these influences, we applied a sliding-window analysis. The window size was set to 450 samples, which is approximately 15 seconds at a 30 Hz sampling rate. This represented the minimum length required to maintain whitened residuals. Windows advanced by 225 samples and the proportion of significant windows per session was quantified to assess whether the relationships were stable or intermittent.

#### Lag-shift and permutation testing

We further explored the alignment of predictive influence by shifting the predictor signal relative to the response across a range of −300 to +300 samples. At each shift, models were refit using the original lag structure. Wald F-tests provided a profile of significance across these offsets to identify where the predictive influence was strongest.

To test for directional asymmetry, we permuted the direction labels across sessions at each lag 10,000 times. This generated a null distribution for the difference in significant session counts. Observed asymmetries were compared to this distribution to compute permutation *p*-values.

### Histological verification

Following the completion of recording sessions, mice were deeply anesthetized with Euthasol and transcardially perfused with phosphate-buffered saline (PBS), followed by 4% paraformaldehyde (PFA) in PBS. Brains were extracted, post-fixed overnight in 4% PFA at 4 °C, and then transferred to 30% sucrose in PBS to prevent prolonged exposure to fixative.

Coronal sections (30 µm) were cut on a vibratome (Leica) through the region containing the 200 µm recording probe track. Sections were collected sequentially into four wells in rotation over 8– 10 rounds, sampling ∼500 µm to include tissue anterior and posterior to the expected probe tip. One full set of sections was mounted and DAPI-stained to screen for the probe track under a fluorescence microscope.

Once the tip was identified, a corresponding section from another well containing the best tip location was processed for immunohistochemistry. These sections were stained for GFP, RFP, and DAPI to verify sensor expression at the recording site. Immunostained sections were scanned with a Zeiss Axioscan.Z1 slide scanner and aligned with DAPI-only sections, providing independent verification of both probe placement and sensor expression.

## Acknowledgements

This research was funded by NIH grant numbers R21AG088596, Brain and Behavior Research Foundation (BBRF) Young Investigator Grant, Whitehall Foundation Research Grant, Welch Foundation Research Grant, Carl B. and Florence E. King Foundation Award, the UT System Rising STARs Award, and UT Southwestern Endowed Scholarship in Medical Science Award.

## Notes

### Competing Interest Statement

The authors have declared no competing interest.

### Summary of Updates

We have added the acknowledgment section to acknowledge the funding sources.

