## Supplemental File for "Coordinated yet asymmetric striatal neuromodulatory dynamics encode associative learning"

Kim et al.

**Includes:**

Supplementary Figures 1 - 6;

Extended Notes;

Supplementary References;

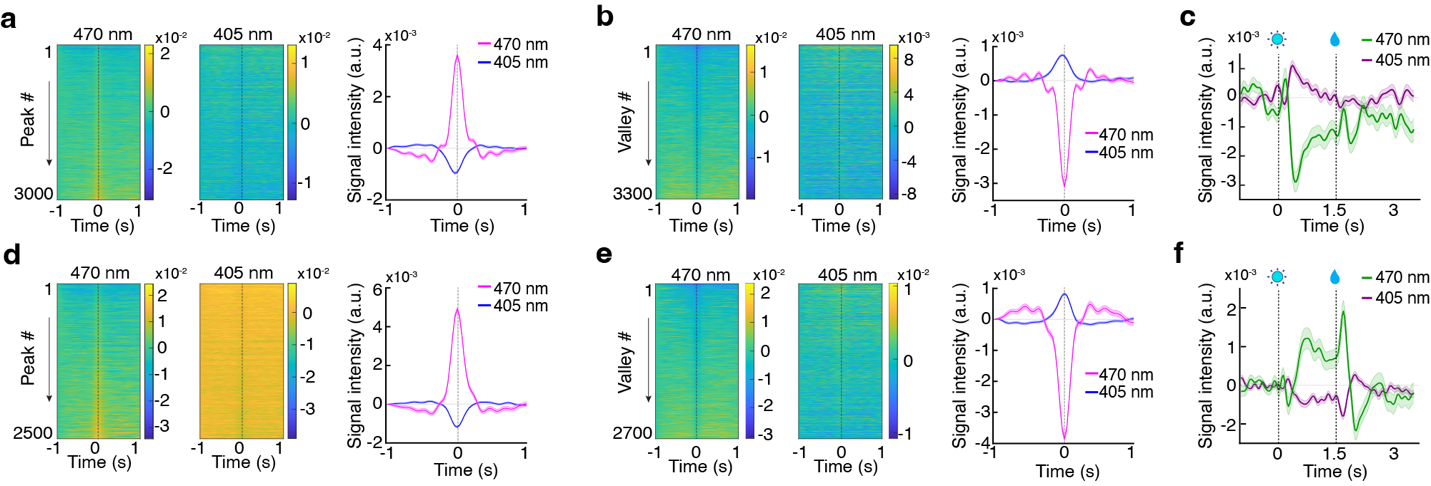

**Supplementary Figure 1 | Inverse modulation between active and isosbestic channels is consistent with hemodynamic coupling rather than motion artifacts.** Data from two representative mice (Animal 1: a–c; Animal 2: d–f) showing the relationship between the active fluorescence signal from the acetylcholine sensor gACh4h (470 nm, excitation) and its isosbestic (ISO, 405 nm, excitation) reference channel. (**a, d**) Peak-triggered analysis: Heatmaps of individual events (left, middle) and mean traces (right) aligned to local maxima in the 470 nm gACh4h signal. (**b, e**) Valley-triggered analysis: Heatmaps of individual events (left, middle) and mean traces (right) aligned to local minima in the 470 nm gACh4h signal. (**c, f**) Task-aligned analysis: Mean raw traces from the late Pavlovian conditioning session, showing concurrent gACh4h (470 nm, green) and ISO (405 nm, violet) signals across cue and reward epochs. Vertical dashed lines indicate cue onset at 0 s and reward delivery at 1.5 s.

**Extended note for Supplementary Figure 1 -** To assess the integrity of our fiber photometry recordings, we performed two complementary validation analyses: (1) characterization of the relationship between the active and isosbestic channels within the green sensor (Fig. S1), and (2) evaluation of cross-channel independence in dual-color recordings (Fig. S2).

*Reciprocal dynamics between active and isosbestic channels.*

Within the green channel, we examined the relationship between the active gACh4h (470 nm) and its isosbestic (405 nm) signals across multiple subjects. Although the 405 nm channel is commonly used to correct motion-related artifacts ^1^, we consistently observed an inverse relationship between the two signals: large fluorescence transients in the active channel were accompanied by time-locked suppressions in the isosbestic signal, whereas troughs in the active signal coincided with increases in the isosbestic channel. While the magnitude of these reciprocal fluctuations was substantially smaller than that of the active-channel response, the effect was consistent and statistically significant.

The observed reciprocal pattern likely arises from a combination of nonexclusive factors, including: (i) wavelength-dependent hemodynamic effects associated with neurovascular coupling. At both 405 nm and 470 nm, oxyhemoglobin exhibits higher absorption than deoxyhemoglobin ^2^, such that increases in local oxyhemoglobin during neural activation could reduce the measured optical signal independently of sensor-specific fluorescence changes ^2,3^. (ii) deviation of 405 nm from the true isosbestic point for the gACh4h sensor, and (iii) local pH fluctuations.

Because these contributions cannot be unambiguously separated in vivo without introducing model-dependent assumptions, we did not explicitly correct for the inverse modulation observed in the 405 nm channel. On this basis, correction with the 405-nm reference channel, as implemented in the preprocessing pipeline, is effective for removing hemodynamic contributions and other nonspecific optical signals. At the same time, because the reference signal may contain additional inverse structure not fully captured by a simple correction model, the standard ΔF/F processing could still introduce a modest amplification of high-amplitude events.

Importantly, this effect was consistent across all subjects, trial types, and learning stages, and does not alter the temporal structure of the signals. Accordingly, our analyses—which focus on temporal dynamics, cross-signal relationships, and condition-dependent contrasts rather than absolute amplitude—are not impacted by this effect. Notably, the opposing dynamics between the active 470 nm and isosbestic 405 nm channels are inconsistent with motion artifacts, which typically manifest as correlated, same-direction fluctuations across both wavelengths.

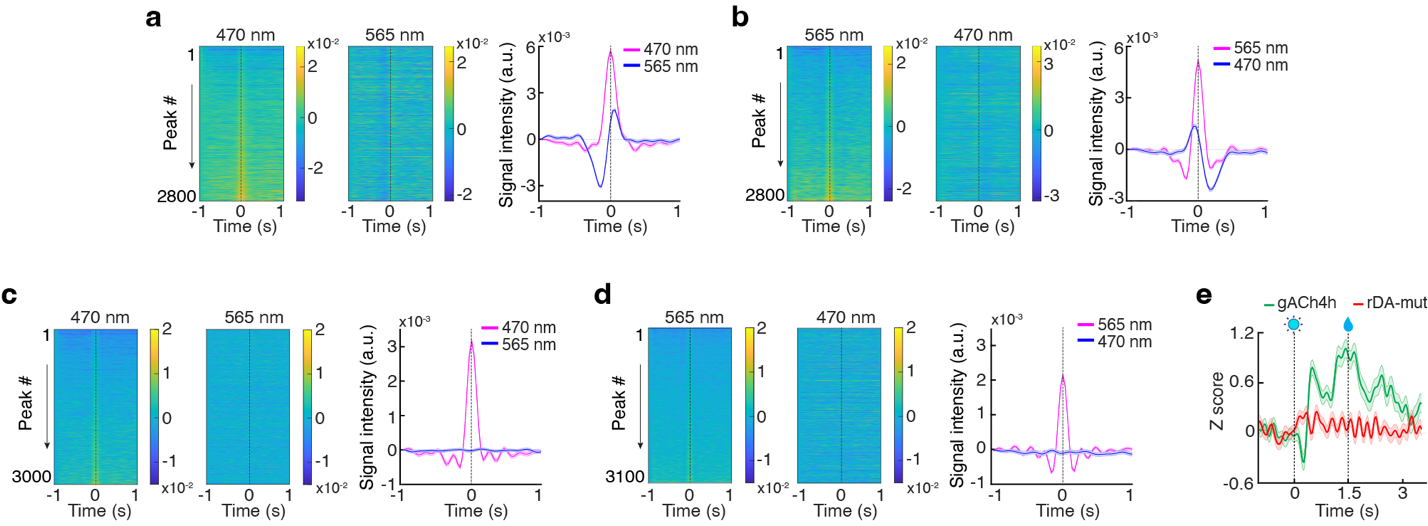

**Supplementary Figure 2 | Mutant-sensor controls and peak-triggered analyses confirm cross-channel independence and neurochemical specificity of dual-color photometry signals.** (**a–b**) Cross-channel peak-triggered analysis of simultaneous acetylcholine sensor (gACh4h) and dopamine sensor (rDA2m) recordings from one representative cue-reward session. Heatmaps of individual events (left, middle) and mean traces (right) show gACh4h signals (470 nm, excitation) and rDA2m signals (565 nm, excitation), aligned to local maxima in the gACh4h channel (**a**) or the rDA2m channel (**b**). **(c–d)** Control recordings from one representative mouse expressing gACh4h and a dopamine–insensitive mutant sensor (rDA-mut). Signals were analyzed the same way as in **a and b,** and aligned to local maxima in the 470 nm gACh4h channel (**c**) or the 565 nm rDA-mut channel (**d**). (**e**) Representative session-averaged dynamics from a late Pavlovian conditioning session (PavL), showing concurrent gACh4h (green) and rDA-mut (red) signals across cue and reward epochs. Mean z-score ΔF/F traces were aligned to cue onset. Vertical dashed lines indicate cue onset at 0 s and reward delivery at 1.5 s.

**Extended note for Supplementary Figure 2 -** To test whether the observed acetylcholine-dopamine (ACh–DA) relationships in dual-color photometry could arise from optical or spectral crosstalk, we performed cross-channel peak-triggered analysis and mutant-sensor controls. In animals expressing functional ACh sensor gACh4h and DA sensor rDA2m, peak-triggered analyses revealed coordinated signal fluctuations across channels, with clearly distinct temporal profiles in 470 nm (gACh4h) and 565 nm (rDA2m) channels (Fig. S2a and b), arguing against trivial bleed-through between channels. We next replaced rDA2m with the dopamine-insensitive mutant sensor rDA-mut ^4^. Under these conditions, peaks detected in the 470 nm channel were not accompanied by corresponding changes in the mutant 565 nm channel (Fig. S2c). Conversely, peaks detected in the mutant channel were not associated with structured responses in the gACh4h signal (Fig. S2d). Consistent with these findings, cue-aligned analysis revealed robust task-related gACh4h rdynamics, whereas the rDA-mut channel exhibited no substantial modulation during cue or reward spochs (Fig. S2e). This stands in clear contrast to the robust 565-nm dynamics observed in mice expressing the functional rDA2m sensor (Fig. 2b and c). Together, these data demonstrate cross-channel independence and indicate that the 470 nm gACh4h signals and the 565 nm rDA2m signals reflect sensor-specific neuromodulatory activity, rather than optical contamination between channels.

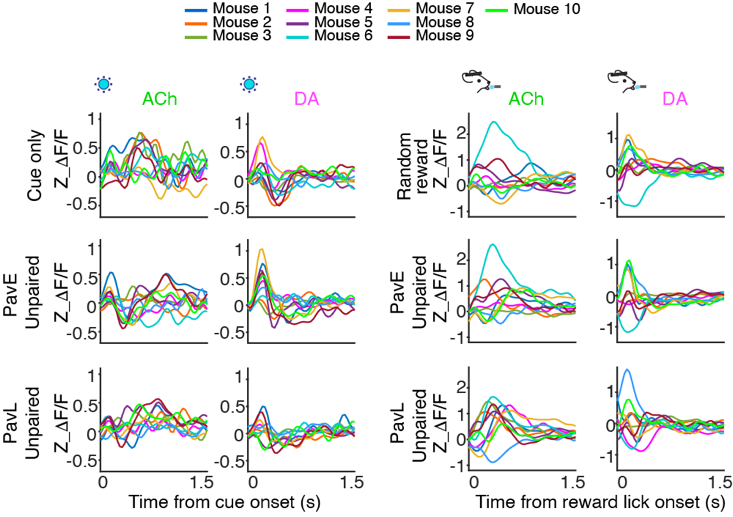

**Supplementary Fig. 3, related to Fig.2. | Unpaired mice show highly variable and session-invariant acetylcholine and dopamine dynamics, lacking the progressive reorganization seen in paired animals.** Population-level dynamics across conditions representing mean ACh and DA traces aligned to cue (left) and reward lick (right) across cue-only, random reward, and Pavlovian training conditions. Colors represent individual mice (n = 10) in the unpaired group.

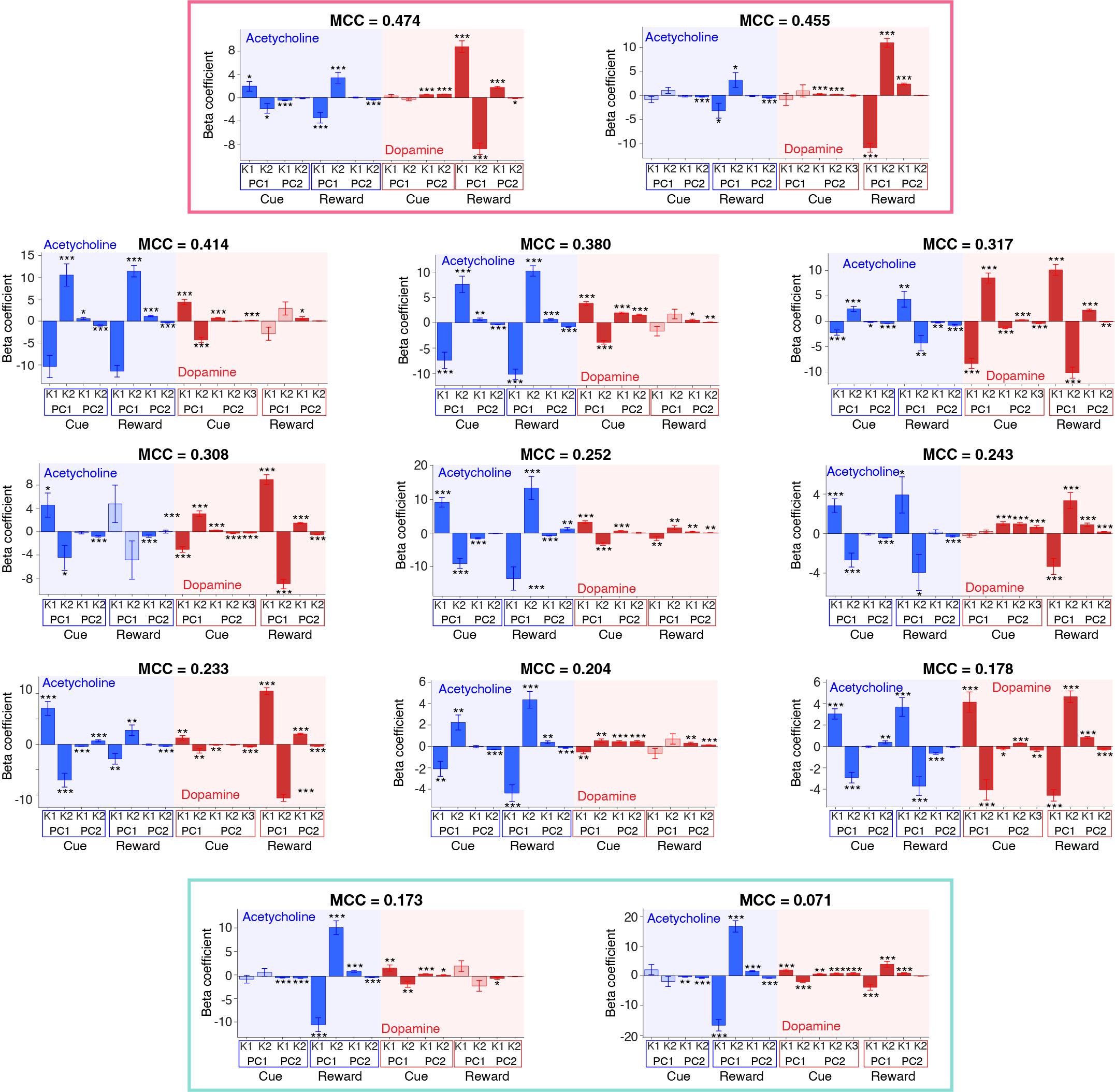

**Supplementary Figure 4, related to Fig. 3 | LOMO fold variability reveals that decoding performance is driven by selective dopamine motif contributions on top of a stable acetylcholine scaffold.** Beta coefficients from individual LOMO decoder folds are ordered by their Matthews correlation coefficient (MCC). Each panel displays motif-specific beta coefficients for acetylcholine (ACh; blue background) and dopamine (DA; red background) across cue- and reward-aligned epochs (PC1–PC2 denote first and second principal components derived motifs; K1–K3 denote individual motif clusters). The top pink box highlights the two highest-performing folds, whereas the bottom green box highlights the two lowest-performing folds. Filled bars indicate statistically significant coefficients (****p* < 0.001, ** *p* < 0.01, * *p* < 0.05), whereas open bars indicate non-significant coefficients. MCC values are shown above each panel.

**Extended note for Supplementary Figure 4**

*Individual decoder performance and neuromodulator contributions.*

Using leave-one-mouse-out (LOMO) cross-validation, we evaluated individual decoder performance with Matthews correlation coefficient (MCC) and examined motif-specific beta coefficients for features derived from acetylcholine (ACh, blue) and dopamine (DA, red) signals, including cue- and reward-aligned components. The analyses revealed distinct contributions of DA and ACh to the learned behavioral state.

Across subjects, ACh motifs, particularly reward-aligned components, consistently exhibited large and significant beta coefficients, consistent with a shared temporal structure across the population. Variation in decoding performance was associated with the degree of DA motif contribution, with decoders with stronger DA weighting tending to achieve higher MCC values. The highest-performing decoders (top pink box) exhibited strong, significant beta coefficients for both the ACh and DA motifs. By contrast, the lowest-performing decoders (bottom green box) retained prominent ACh reward-related weights but showed weaker DA contributions.

Together, these results suggest a structured division of roles within the low-dimensional motif space, in which ACh reflects a stable population-level component, whereas DA provides a more selective feature that contributes to accurate decoding of the learned state.

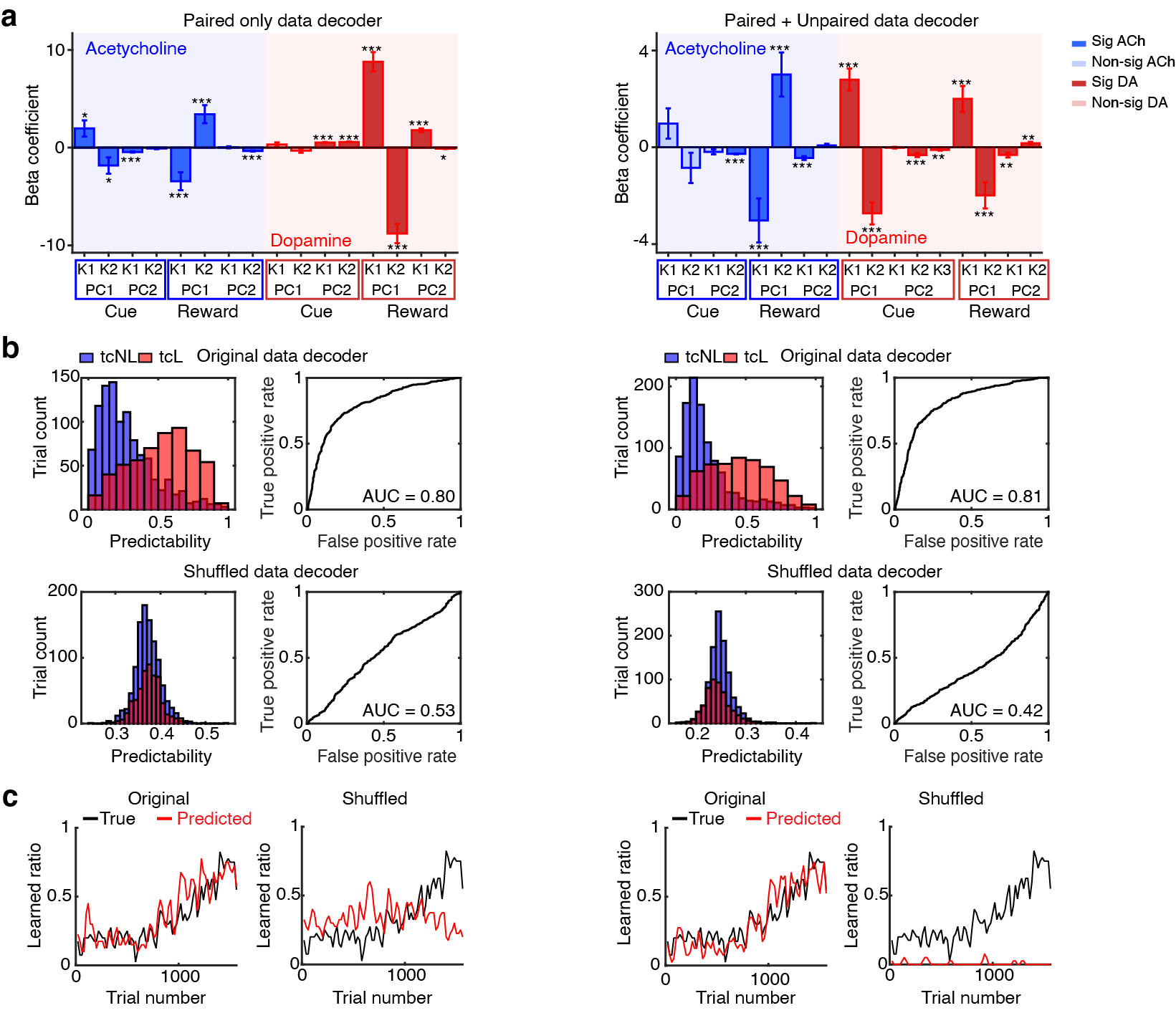

**Supplementary Figure 5, related to Fig. 3 | Consistent decoding performance and motif geometry across training datasets reinforce the functional dissociation between acetylcholine and dopamine representations.** (**a**) Comparison of beta coefficients between LOMO decoders trained on Paired group-only data (left) and on the full dataset including both Paired and Unpaired groups (right; ****p* < 0.001, ** *p* < 0.01, * *p* < 0.05). (**b**) Decoding performance for original (top) and shuffled-label (bottom) models. Both LOMO decoders show robust classification performance on the original data (AUC ≈ 0.81), whereas shuffled-label controls exhibit near-chance performance. (**c**) Comparison of true (black) and predicted (red) proportions of learned trials across trials.

**Extended note for Supplementary Figure 5**

To evaluate the generalizability of the decoding framework and its specificity for associative learning–related structure, we performed a leave-one-mouse-out (LOMO) analysis on a representative high-performing subject (Mouse #1), comparing decoders trained on two data scales: (i) the Paired-only group and (ii) the full dataset, including both Paired and Unpaired groups.

*Preservation of motif structure across training scales (Fig. S5a).*

Both training regimes recovered highly similar low-dimensional motif structure. Beta coefficients associated with ACh and DA principal component motifs showed broadly conserved weighting patterns across cue- and reward-related epochs, indicating that expanding the training set with unpaired animals did not substantially alter the learned decoder structure. Rather, motif contributions were preserved across training compositions, supporting the robustness of the low-dimensional representation.

*Decoder performance and shuffle controls (Fig. S5b and c).*

Both decoders accurately tracked the learning trajectory of Mouse #1 and exhibited comparable performance (AUC ≈ 0.80–0.81), demonstrating cross-animal generalization stable across training scales (Fig. S5b upper panels). To test whether this performance could arise from spurious signal structure, we trained shuffle-control decoders using permuted labels. In both cases, shuffled models collapsed to near-chance performance (AUC ≈ 0.42–0.53; Fig. S5b bottom panels), with predicted probabilities failing to separate learned from unlearned trials and failing to track the behavioral transition (Fig. S5c). These controls indicate that decoder performance depends on genuine neural–behavioral structure rather than incidental statistical regularities.

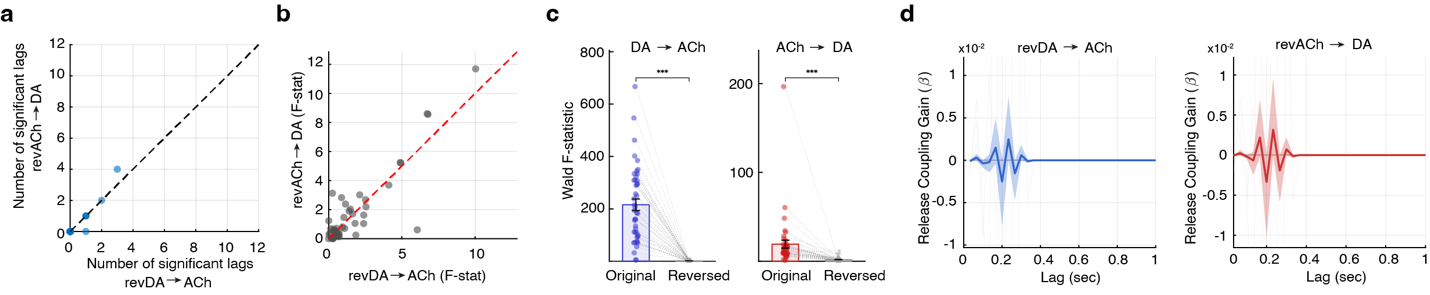

**Supplementary Figure 6, related to Fig.5 | Time-reversal abolishes the directional asymmetry of Granger causality, confirming a temporally specific biological basis for dopamine-to-acetylcholine influence.** (**a**) Comparison of significant lag numbers for the reciprocal GC directions in time-reversed data: reversed **DA → ACh** versus reversed **ACh → DA**. (**b**) Scatter plot of Wald F-statistics computed from time-reversed data for the two GC directions. (**c**) Group comparisons of Wald F-statistics for original (colored dots) versus time-reversed (gray dots) data for the DA → ACh model (left; ****p* < 0.001) and the ACh → DA model (right; ****p* < 0.001). (**d**) Lag-resolved impulse response functions (gain β) across 0–1 s for the time-reversed analysis, shown for reversed DA → ACh in blue (left) and reversed ACh → DA in red (right). Shaded regions represent 95% bootstrap confidence intervals.

**Extended note for Supplementary Figure 6**

To assess whether the observed DA → ACh directional influence reflects a true temporal structure rather than intrinsic autocorrelation or shared signal properties, we performed a time-reversal control analysis using data from 22 mice (n = 12 Paired, n = 10 Unpaired; first and last Pavlovian sessions). In contrast to the forward-time analysis (Fig. 5b, c), time-reversed data exhibited a marked reduction in significant lags and no consistent directional bias (Fig. S6a, b). The large F-statistics observed in the forward-time analysis (peak F ≈ 800; Fig. 5c) are strongly attenuated following time reversal (peak F < 14; Fig. S6c), and the directional asymmetry favoring DA → ACh is no longer present. Direct group comparisons further confirmed this effect, showing a significant loss of predictive gain in both DA → ACh and ACh → DA models upon time reversal (*** *p* < 0.001, Wilcoxon signed-rank test). Consistent with these results, the temporal profile of the interaction, quantified by the coupling gain (β), exhibited a distinct positive peak in the DA → ACh direction at lags of 0.2–0.4 s in the forward-time data (Fig. 5d). This structured lag-dependent coupling was absent in the time-reversed analysis, with β coefficients reduced to near-baseline levels across all lags (Fig. S6d). Together, these findings indicate that the observed directional interaction reflects a temporally specific biological process rather than an artifact of shared autocorrelation structure.
